# MicroFisher: Fungal taxonomic classification for metatranscriptomic and metagenomic data using multiple short hypervariable markers

**DOI:** 10.1101/2024.01.20.576350

**Authors:** Haihua Wang, Steven Wu, Kaile Zhang, Ko-Hsuan Chen, Rytas Vilgalys, Hui-Ling Liao

**Affiliations:** North Florida Research and Education Center, University of Florida, 155 Research Road, Quincy, FL, USA; Department of Soil, Water, and Ecosystem Sciences, University of Florida, Gainesville, FL, USA; Departement of Agronomy, National Taiwan University, Taipei, Taiwan; Biodiversity Research Center, Academia Sinica, 128 Academia Rd., Nangkang Dist., Taipei, Taiwan; Department of Biology, Duke University, 130 Science Drive, Durham, NC, USA

**Keywords:** MicroFisher, fungal classification, hypervariable marker databases, metagenomics, metatranscriptomics

## Abstract

Profiling the taxonomic and functional composition of microbes using metagenomic (MG) and metatranscriptomic (MT) sequencing is advancing our understanding of microbial functions. However, the sensitivity and accuracy of microbial classification using genome– or core protein-based approaches, especially the classification of eukaryotic organisms, is limited by the availability of genomes and the resolution of sequence databases. To address this, we propose the MicroFisher, a novel approach that applies multiple hypervariable marker genes to profile fungal communities from MGs and MTs. This approach utilizes the hypervariable regions of ITS and large subunit (LSU) rRNA genes for fungal identification with high sensitivity and resolution. Simultaneously, we propose a computational pipeline (MicroFisher) to optimize and integrate the results from classifications using multiple hypervariable markers. To test the performance of our method, we applied MicroFisher to the synthetic community profiling and found high performance in fungal prediction and abundance estimation. In addition, we also used MGs from forest soil and MTs of root eukaryotic microbes to test our method and the results showed that MicroFisher provided more accurate profiling of environmental microbiomes compared to other classification tools. Overall, MicroFisher serves as a novel pipeline for classification of fungal communities from MGs and MTs.

## Background

Microbiota that comprises the majority community in terrestrial and marine ecosystems as well as within an organism (such as animals and plants) perform crucial roles in regulating biochemical and biological mechanisms in these systems. Therefore, it is critical to study the community and functional composition of microorganisms to understand whole system biology and ecosystem functions[1]. Next-generation sequencing approaches, including metagenomics (MGs) and metatranscriptomics (MTs), provide revolutionary ways to identify critical functions of microorganisms in environments and within an host organism[2–5]. In addition to functional profiling, MGs and MTs offer opportunities to characterize microbial community composition and diversity at multigene scales[6]. However, the classification of taxonomy through MG and MT sequencing remains challenging due to the limitations of current sequencing platforms (e.g., short reads) and the lack of comprehensive genome databases.

The current methods for taxonomic profiling using MG and MT sequences mainly rely on the read-mapping approach[7–9] and the read-assembly approach[10–12]. The read– (or k-mer) mapping approach, as used by tools such as Kaiju[8], Kraken2[7], and Metaphlan3[9], is a method for profiling the microbial taxonomy via mapping the reads to pangenomic references, followed by providing taxa prediction. However, this approach relies on the availability of genome references, which limits the identification of most taxa in environmental samples. For instance, using a fungal genome database containing 949 fungal species genomes (∼32 Gb) to study pig and mouse microbiomes, only 71 fungal species were detected[13]. This indicates that the limitation in the available genome references significantly reduced the resolution of fungal profiling in MG and MT datasets[14,15]. By contrast, the read-assembly approach allows for the identification of the microbial community composition via performing the de-novo assembly and binning to reconstruct genomes and identify the taxonomic composition of bacteria, archaea, and viruses[16,17]. However, it is challenging to generate accurate *de-novo* assembly results for eukaryotes from MG and MT using this approach, due to the large genome size, highly conserved regions, and repetitive regions across some taxa present in eukaryotic genomes that lead to a high number of false positives mixed with true identifications[18]. The limitations of these two methods impede the comprehensive understanding of metagenomic-level community composition and function present in complex ecosystem systems, particularly in eukaryotes. Thus, it is crucial to develop a non-genome but multi-gene/sequence region-based classifier to improve the accuracy and precision of fungal classification in shotgun sequencing datasets.

Although the pangenome-[7,9] and core protein domain-based[19] profiling pipelines have been widely used in the classification of MG and MT sequencing, rRNA (such as SSU and LSU) and ITS markers have been recognized to more accurately identify the taxonomic composition and provide a more reliable prediction of microbial communities[20]. This is because these marker genes provide taxa-specific fingerprints for short-read classification[21]. It has a long history of identifying fungal taxa based on the marker genes from ribosomal RNA (rRNA) regions. Until now, marker genes have been widely used in fungal identification, such as the small subunit (SSU) rRNA, internal transcribed spacer (ITS), and large subunit (LSU) rRNA regions are popularly applied in microbial community profiling of DNA amplicon sequencing. The elongation factor[22] and beta-tubulin[23] have also been used to identify certain taxonomy groups. Over the years, researchers have developed a variety of analysis tools and many databases which contain a massive number of marker sequences for fungal taxonomic classification, such as UNITE[24], SILVA[25], and RDP[26]. However, the rRNA and ITS marker sequences cannot be directly used in either read-assembly or read-mapping methods for the short reads, due to the highly conserved regions present in these marker sequences that can cause a fatal bias[5]. Thus, we applied hypervariable regions in rRNA and ITS sequences to classify fungal taxa.

We present MicroFisher, a computational workflow for fungal classification that can produce accurate and sensitive taxonomic profiles. We generated hypervariable marker databases (HMDs) consisting of several highly variable regions of LSU rRNA and ITS marker sequences, which can provide species-specific fingerprints for fungal short-reads identification. Briefly, MicroFisher first assigns the fungal taxonomic composition for an individual MG or MT sample by mapping the intact short reads onto these HMDs. The taxa abundances are estimated from multiple databases and used a “weight-based abundance integration algorithm” to generate the relative taxonomic abundance for profiling the fungal community from MTs and MGs. Overall, this study provides the first-step guide for accurately profiling the fungal community from short-read datasets by applying hypervariable markers. In the long run, the MicroFisher protocol is capable of incorporating newly developed marker sequences that are specialized for certain fungal groups.

## Methods

### (A) Generation of hypervariable marker databases (HMDs)

In this study, we selected four sets of hypervariable marker sequences, including LSU rRNA D1 and D2 domains as well as ITS1 and ITS2, which represent highly variable marker sequences for fungal identification. We further identified and selected higher polymorphic sub-regions among these marker sequences to serve as hypervariable marker databases (HMDs) (Fig. 1A).

**Fig. 1.**
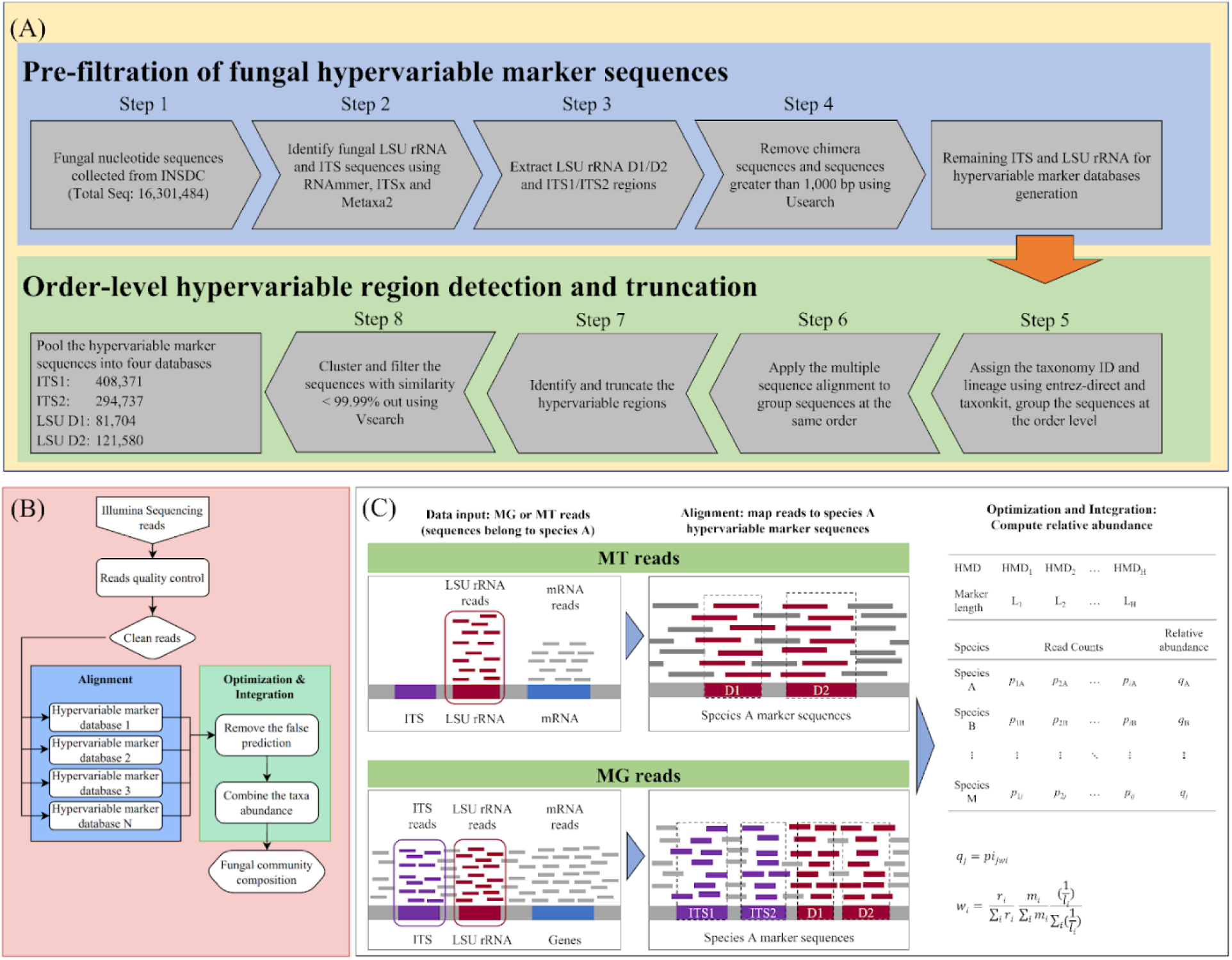
Overview of hypervariable marker database generation and the principle of MicroFisher classification. (A) Computational flowchart for the generation of hypervariable marker databases. (B and C) MicroFisher pipeline for fungal identification and abundance estimation in metagenome (MG) and metatranscriptome (MT) analysis (1) Alignment: aligning the short reads in MG and MT to multiple hypervariable marker databases; (2) Optimization and integration: use a weight-based algorithm to remove false positives and optimize the estimated abundance of each taxon from multiple hypervariable databases. The final output results were returned fungal community composition across taxonomic levels. *q_ij_* is the weighted proportion for an identified taxon *j* against HMD *i*, *p_ij_* is the original abundance of identified taxa *j* estimated by Centrifuge against HMD *i*, and *w_i_* is the weighting associated with the abundance report from a particular HMD *i*. *r_i_* is the total number of reads obtained in the Centrifuge classification aligned to HMD *i*, *m_i_* is the minimum length parameter (MiniHit) used in the Centrifuge classification for the reads aligned to HMD *i*, and *l_i_* is the mean length of hypervariable marker sequences in HMD *i*.

#### (a1) Pre-filtration of fungal hypervariable marker sequences (Fig. 1A, Steps 1 to 4)

We developed a four-step process to pre-filter the marker sequences from the fungal nucleotide sequences originally obtained from International Nucleotide Sequence Database Collaboration (INSDC)[27]. Specifically, a total of 16,301,484 fungal nucleotide sequences, including genomic DNA, mRNA, and rRNA, were collected (Step 1). In Step 2, fungal rRNA sequences were predicted by a hidden Markov model with RNAmmer[28] and HMMER2[29], and non-LSU rRNA sequences were rejected using Metaxa2[30] with default parameters. In Step 3, fungal LSU rRNA sequences were extracted: a custom pipeline based on the fungal LSU rRNA universal primer sequences was subsequently applied to search the LSU rRNA D1 and D2 regions using SeqKit[31]. A similar workflow was applied to fungal ITS sequences. Briefly, after filtering the non-ITS sequences using a fungal ITS universal primers-based custom pipeline, ITSx[32] was applied to predict the fungal ITS region with ITS1, 5.8S rRNA, and ITS2. Chimeric sequences in the ITS database were then detected and removed using UCHIME (package: *Usearch*)[33] in high confidence mode. The extracted ITS and LSU rRNA D1 and D2 sequences served as input for the next step after removing sequences with unexpected lengths (sequence length > 1000 nt) (Step 4).

#### (a2) Detection of hypervariable marker region (Fig 1A, Steps 5 to 8)

We applied Entrez-direct[34] and Taxonkit[35] to assign the taxonomy to each sequence collected from Step 4 against the NCBI taxonomy database (Step 5). Sequences assigned to the same order were grouped together, and a multiple sequence alignment was performed for each group using the Clustalw package[36] with default parameters (Step 6). The multiple sequence alignment results were used to identify the “hypervariable regions”. The regions with low consensus (containing more gaps) were candidate regions for “hypervariable regions” (Step 7) (Fig. S1). Each base site was quantified among these candidate regions using the bit value that was calculated based upon the equation described by Bembom (2007)[37]. Using seqLogo, the regions with low bit values between conserved flanking regions (high bit values) were identified as the “hypervariable regions” to serve as marker fingerprints for fungal classification[38],[39] (Fig. S2). Using this approach, general “hypervariable regions” were identified. Then, the conserved flanking regions of “hypervariable regions” for a sequence were truncated and the remaining subset of hypervariable regions was clustered and deduplicated to remove the duplicate sequences with similarity > 99.99% by Vsearch package[40] (Step 8). The hypervariable sequences from different taxonomic orders were then merged into a corresponding single marker sequence, and four hypervariable marker databases ITS1, ITS2, LSU D1, and D2 were curated accordingly. The average length of each marker is 188.8 bp for ITS1, 186.8 bp for ITS2, 172.8 bp for D1, and 189.4 bp for D2, respectively (Table S1).

In this study, HMDs generated include 405,672 sequences in ITS1 database, 294,737 sequences in ITS2 database, 81,704 sequences in LSU D1 database, and 121,580 sequences in LSU D2 database (Table S1). A total of 903,693 hypervariable marker sequences from these four HMDs span 10 phyla, 62 classes, 234 orders, 873 families, 6,545 genera, and 104,072 species. Around 19,714 fungal species were shared in all four databases (Fig. S3). Since the process of creating these short HMDs is computationally intensive, we provided the latest reference databases and updated information for the users to download. Users can directly use these HMDs in the MicroFisher pipeline.

### **(B)** MicroFisher pipeline: classification, integration, and optimization

We developed the MicroFisher pipeline to identify fungal taxa and estimate their relative abundance from the qualified MG and MT sequences (Fig. 1B).

#### (b1) Alignment: Align the qualified reads of MG and MT to HMDs

After quality trimming, the MGs and MTs reads generated from the environmental samples (e.g., soils, marine, plants, and animals) served as the input files for the MicroFisher pipeline. Firstly, the MicroFisher classifier searches these reads against selected HMDs. The Centrifuge [41] was used by MicroFisher as the microbial classification engine which identifies the potential taxa and estimates taxa abundance. In this step, the minimum hit (MiniHit) length is one of the key parameters in Centrifuge to determine the threshold for reads matching a marker sequence in HMDs. Different from Centrifuge, which only applies a single HMD during classification, MicroFisher integrates searching reports from Centrifuge using multiple HMDs for microbial classification. Therefore, the MiniHit length is critical for the classification sensitivity and accuracy of MicroFisher.

#### (b2) Integration and optimization: Integrate abundance reports using weight-based algorithms

The MicroFisher pipeline collects and integrates the abundance reports obtained from the reads aligned to different HMDs using weight-based algorithms and generates a “final relative abundance table”. The application of weight-based algorithms improves the accuracy of taxonomic classification and abundance estimation (Fig. 1C).

Three determined factors are incorporated in the algorithms to weight the abundance reports: 1) the total number of reads aligned to each HMD (e.g., more reads aligned to the HMD, the higher weight), 2) the MiniHit length, and 3) inverse weighting of the average length of hypervariable marker sequences in each HMD (i.e., Longer the average length, the lower the weight) (as recorded in Table S1). Incorporating these factors in the algorithms improves both the accuracy and precision of fungal taxonomic identification.

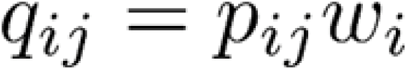

Where *q_ij_* is the weighted proportion for an identified taxon *j* against HMD *i*, *p_ij_* is the original abundance of identified taxa *j* estimated by Centrifuge against HMD *i*, and *w_i_* is the weighting associated with the abundance report from a particular HMD *i*, which is subsequently used to merge reports from multiple HMDs. *w_i_* can be calculated by the following formula:

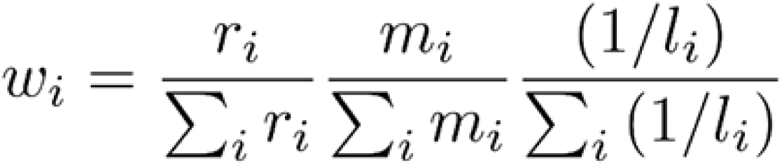

Where *r_i_* is the total number of reads obtained in the Centrifuge classification aligned to HMD *i*, *m_i_* is the minimum length parameter (MiniHit) used in the Centrifuge classification for the reads aligned to HMD *i*, and *l_i_* is the mean length of hypervariable marker sequences in HMD *i*.

The value of these three determined factors (number of reads, MiniHit length, average length of HMD) can be configured by the users to achieve alternative weighting schemes for the taxonomic classification results combination if needed. By setting all parameters to the same value for a dataset aligned to different HMDs, MicroFisher may apply equal weighting for that dataset across all HMDs.

### **(C)** Using MicroFisher to profile fungal community with synthetic and real data

#### (c1) Fungal community profiling for synthetic data

We evaluated Microfisher performance in fungal community profiling using synthetic MG data. Synthetic MG data [NovaSeq (151 bp)] were generated by InSilicoSeq to serve as the synthetic data input[42]. The fungal rRNA sequences were obtained from the NCBI RefSeq database. We simulated three fungal taxa pools by randomly selecting 50, 100, and 200 fungal species from 6,886 species with their sequences covering both ITS and LSU rRNA regions (Table S2). Five synthetic fungal communities were generated for each pool level, resulting in a total of 15 synthetic communities [5 million (M) noisy reads in each sample]. The reads of the selected fungal species in these synthetic communities were distributed exponentially to represent the gradient of dominance toward rare fungal taxa. These 15 synthetic community datasets then served as the data input for fungal classification.

Four HMDs generated in section (a), including ITS1, ITS2, LSU D1, and LSU D2 were used for single HMD classification with Centrifuge. Meanwhile, the MicroFisher package was applied with the default algorithm described in section (b) using three HMDs combinations, including ITS1 + ITS2 HMDs (ITS only: ITS1+ITS2), LSU D1 + LSU D2 HMDs (LSU only: LSU D1+D2), and a combination of four HMDs (ITS+LSU). We evaluated the effects of the MiniHit length parameter (between 70-150 bp) on the prediction of the fungal community profile. The accuracy of abundance tables generated from Centrifuge and MicroFisher was evaluated by comparing the original synthetic fungal species pools.

Three metrics were applied to evaluate the performance of MicroFisher in fungal classification, including accuracy, precision, and sensitivity (Fig. 2). The true positive (TP), false positive (FP), and false negative (FN) of fungal taxa classification across different taxonomic levels were identified. The calculations of these metrics followed the formula described by Lu and Salzberg (2020)[5]. Briefly, the sensitivity was defined as TP/(TP+FN); the accuracy was calculated by TP/(TP+TN+FN), and precision was calculated by TP/(TP+TN) (Definition see Table S4). The metrics, including sensitivity, accuracy, and precision, were plotted to illustrate the effect of the combination algorithm, MiniHit length, and community complexity on MicroFisher’s prediction performance (Fig. 2 and Fig. S4).

**Fig. 2.**
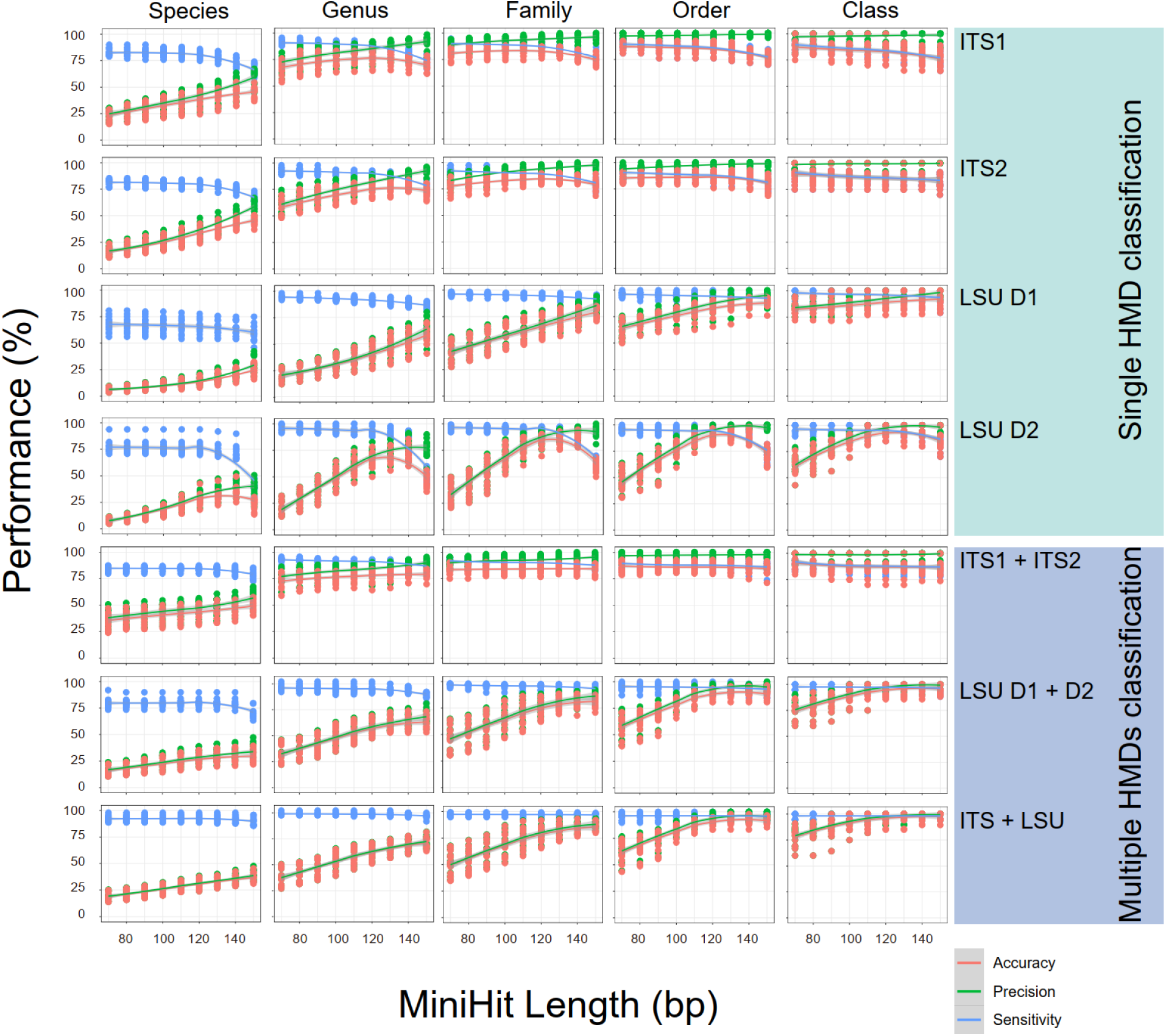
The performance of MicroFisher’s fungal classification. The fungal classification and profiling were performed on these mock communities using MicroFisher (multiple HMDs applied) and Centrifuge (single HMD applied). The synthetic communities (151 bp) were simulated from three levels of fungal composition complexity (50, 100, and 200 fungal species), with five synthetic communities for each level. The performance of accuracy (in red), precision (in green) and sensitivity (in blue) were generated for each taxonomy level using Minimum Hit Length (MiniHit Length) range from 70 – 150 bp.

Furthermore, the histogram plots of predicted abundance were applied to compare the relative abundance distribution of true and false positives (Fig. 3A and Fig. 4). We also calculated the absolute error and root mean square (r.m.s.) error of the relative abundance tables generated from MicroFisher (Figs. 3B and S5). The absolute error of a fungal taxon indicates the difference of estimated relative abundance subtracted from actual relative abundance. The r.m.s error was further calculated to measure the error rate of taxa abundance estimates (formula listed in Table S4). The Pearson correlation analysis of estimates and actual relative abundance of taxa was performed (Fig. 3C and Figs. S6-S8). The statistical analysis for Fig. 2 and 3 was conducted with R (version 4.1.3).

**Fig. 3.**
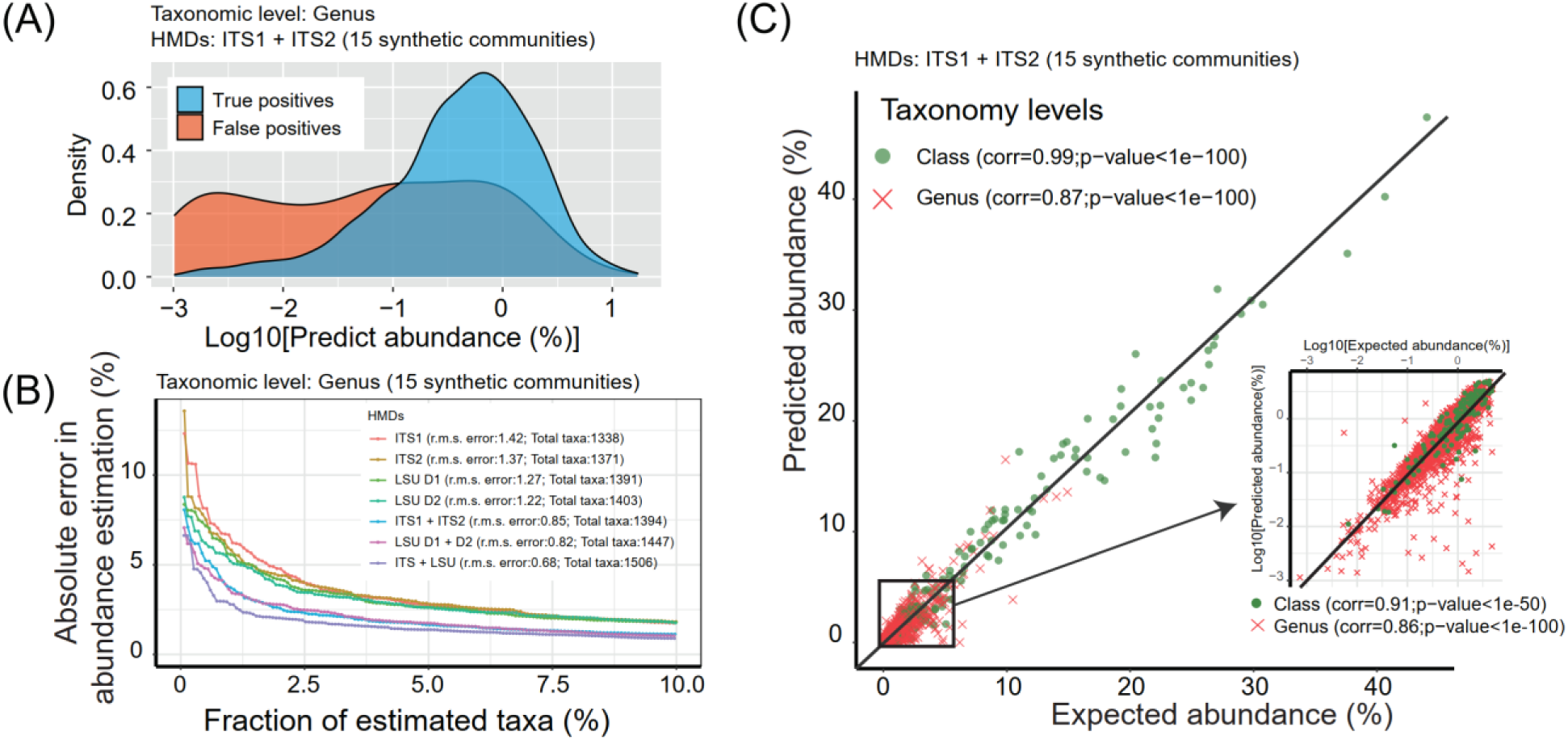
Evaluation of Accuracy of relative abundance tables generated from 15 synthetic fungal community data using MicroFisher. Fungal classification and abundance estimation were performed using a MiniHit length of 120 bp in the MicroFisher pipeline. (A) Density plot shows the patterns of the fungal abundance (genus level) between true positives and false positives. (B) Absolute and root mean square (r.m.s) errors estimated while predicting the abundance of fungal taxa at genus level using single HMD (ITS1, ITS2, LSU D1, and LSU D2; profiled by Centrifuge with minimum hit length of 120 bp) and multiple HMDs (ITS1+ITS2, LSU D1+D2, and ITS+LSU; profiled by MicroFisher with default setting). The taxa were ordered by the value of absolute error and exhibited with a fraction of the estimated taxa. Root mean square (r.m.s.) error was calculated for each HMD combination, and the total number of true positive clades detected by different modes from 15 synthetic communities was indicated as “Total taxa: numbers”. (C) Correlations of true and inferred taxa abundances for taxa estimated from these 15 synthetic communities. Green and red dots denote taxa at the class and genus level respectively. To explore the correlation of taxa with abundance less than 5 %, we transformed them using log10 before performing the Pearson correlation. Pearson correlation coefficient and P-value for the taxa were calculated. MicroFisher classification and abundance evaluation were conducted using ITS1+ITS2 HMDs with default parameters (MiniHit length of 120 bp, weighted and filter mode).

**Fig. 4.**
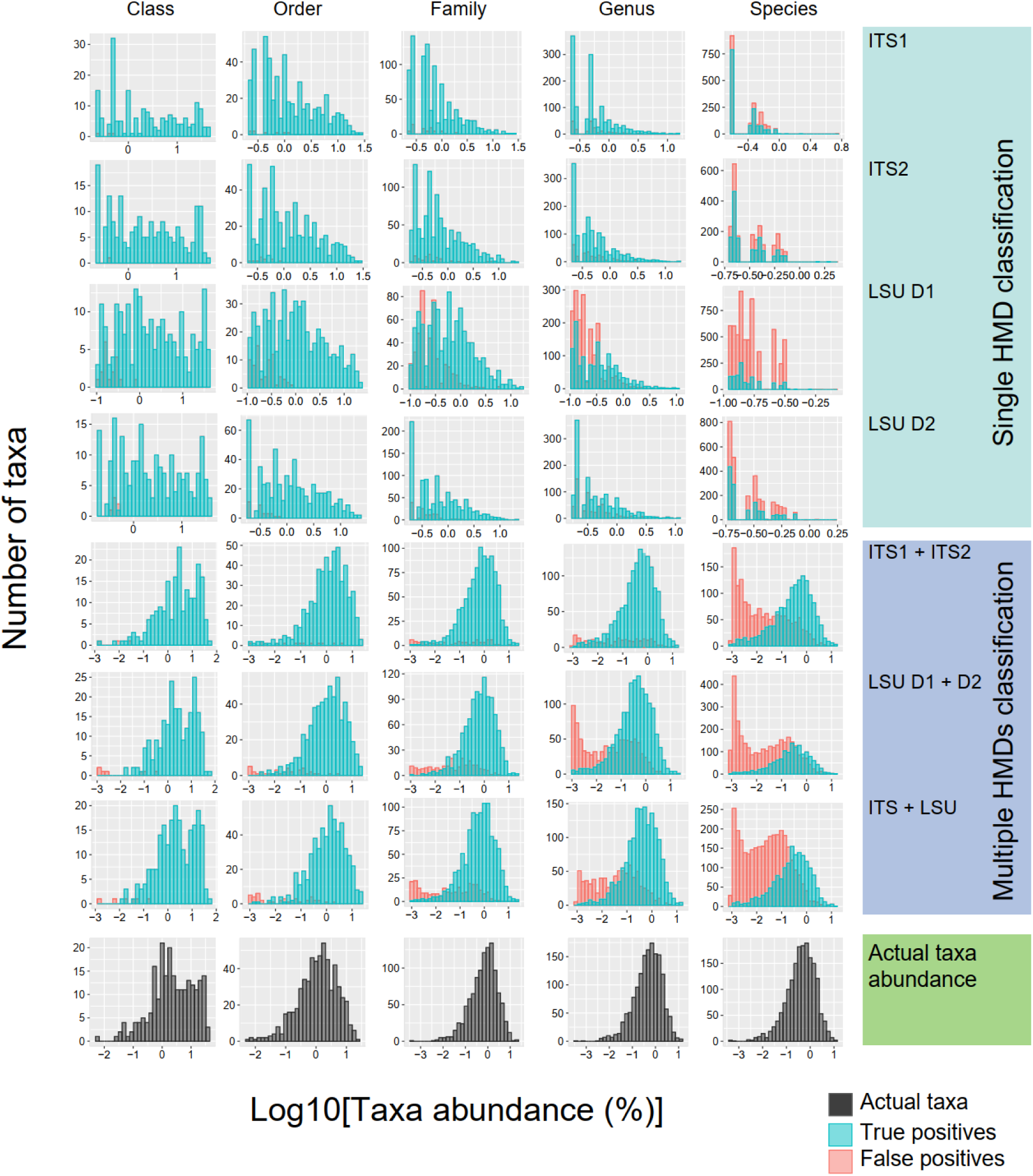
Histogram plots compare the patterns of the fungal abundance (genus level) between predicted results and actual data. The single HMDs (ITS1, ITS2, LSU D1, and LSU D2; profiled by Centrifuge) and multiple (ITS1+ITS2, LSU D1+D2, and ITS+LSU; profiled by MicroFisher with default setting) were applied to compare the similarity of taxa abundance patterns. The minimum hit (MiniHit) length of 120 bp was applied for Centrifuge and MicroFisher.

#### (c2) Fungal community profiling from real MG and MT data

We further applied MicroFisher to classify fungal composition and estimate its abundance using real MG and MT data (Table S3). To compare the performance of MicroFisher with other alignment-based taxonomic classification techniques, we also ran MG and MT datasets across other classification tools with default settings, including Eukdetect, Kaiju, Kraken2, and Metaphlan3. The amplicon sequences from the same samples that were used to generate MG data were also analyzed in order to compare the fungal classification performance of MicroFisher or other tools from MGs to standard amplicon sequencing. The data input selected for this study includes pine forest soil MGs, their amplicon sequences, and pine root MTs. The soils were sampled from *Pinus* forests located in Jelonan and Belanglo in Australia and Cambria, Monterey, and Swanton in the United States. Total soil DNA was extracted using the DNA PowerSoil kit (MoBio, Carlsbad, California, USA) following the manufacturer’s instructions. A two-step PCR approach[43] targeting ITS1 and ITS2 regions was used to generate the DNA library, and the mixture of amplicons was sequenced on an Illumina (Illumina Inc., San Diego, CA, USA) Miseq (v3 300 bp, 13 Gb sequencing capacity) at the Duke Center for Genomic and Computational Biology, United States. Shotgun metagenomic sequencing of the same soil DNA samples was performed at DOE Joint Genome Institute (JGI), United States. Moreover, the collected soils were used to grow *Pinus radiata* seedlings for three months and the pine roots were harvested. Total RNA was extracted from the collected pine root for metatranscriptomic sequencing at JGI. Soil fungal ITS amplicon sequences were processed using QIIME2[44], and the bioinformatic details can be found in Zhang et al. (2022)[45]. The community compositions of soil metagenomics, root metatranscriptomics, and respective metataxonomics were compared and visualized using the ggplot2[43],[46] package in R.

## Results

### Database and pipeline overview

We compiled the HMDs targeting the hypervariable regions of fungal ITS, LSU D1 and D2, containing 104,072 fungal isolates within 6,545 genera (Fig. 1A, Table S1). The HMDs provide high-resolution (by applying hypervariable sequence regions) and multiple set references, which reduces the number of mismatches (false positives) and false negatives in fungal classification. The trimming steps (Steps 6-7 in Fig. 1A) based upon sequence consensus allow us to keep only the hypervariable region for HMDs, which improves the accuracy of fungal taxa predictions. The flanking regions of the trimmed sequences in HMDs are less than 10 bp to minimize the false positives in the fungal classification. The marker sequences have a wide range of lengths (120 – 350 bp) to enhance variability.

We developed the MicroFisher package, which employs multiple HMDs (including ITS1, ITS2, LSU D1, and LSU D2) to classify fungal composition and estimate taxa abundance from MG and MT sequencing (Figs. 1B and 1C). The first step of MicroFisher pipeline is to map reads to several HMDs and generate corresponding abundance reports using Centrifuge. A weight-based algorithm is further employed to optimize and integrate those Centrifuge reports into a final abundance table of detected taxa. The weight-based integration algorithm takes into account the total number of mapped reads, MiniHit length, and average sequence length of the mapped data. Thus, the weight-based integration algorithm effectively reduces mismatches (false positives), and the combination of multiple HMDs reduces the chance of missing taxa (false negatives).

### Fungal community profiling from a synthetic community

We applied the MicroFisher pipeline on a synthetic metagenomic dataset to examine the performance of MicroFisher in fungal classification and abundance estimation. Overall, most species from synthetic metagenomic datasets (∼92%) were identified using MicroFisher. The results showed that the combination of HMDs usage and MiniHit length of the mapped reads primarily determined MicroFisher’s performance. For example, fungal community classification performance was improved when multiple HMDs were used instead of a single HMD (Fig. 2). By applying MiniHit at 120 bp for genus-level detection, the average sensitivity, accuracy, and precision using multiple HMDs vs. single HMD were 95%, 66%, 69%, vs. 90%, 64%, 69%, respectively. At the genus level of MiniHit 120 bp, using LSU D1+D2 HMDs resulted in 95% sensitivity, 59% precision, and 57% accuracy of performance, while applying LSU D1 HMD only gave 91% sensitivity and 42% precision, and 40% accuracy. In addition, we found that the prediction accuracy and precision of MicroFisher increased with increasing MiniHit length (Fig. 2). Compared with MiniHit lengths of 80 bp and 100 bp, setting MiniHit length at and higher than 120 bp gave better accuracy, precision, and sensitivity. On average, 36%, 69%, and 83% precision of taxa prediction were detected across species, genus, and family levels at a MiniHit length of 120 bp, respectively. Reduction of MiniHit length to 80 bp reduced the precision to 27%, 53%, and 66% across these taxonomic levels (summarized result from Fig. 2). At MiniHit length of 120 bp, 86%, 95 %, and 94% of fungal taxa on average were recovered across species, genus, and family levels, respectively, reflecting high sensitivity in fungal prediction using MicroFisher. However, MiniHit of 150 bp only recovered 79%, 91%, and 92% of expected fungal taxa. To obtain an optimal classification performance, the MiniHit length of 120 nt was then set as the default parameter in the MicroFisher pipeline. More importantly, the genus and higher taxonomic levels were recommended for subsequent analysis in our pipeline because of their high precision values (> 60%).

We further evaluated the accuracy of relative abundance tables generated from MicroFisher (Fig. 3). After applying the weight-based integration algorithm, a similar pattern was observed between predicted abundance with true positives and actual abundance, especially when a multiple HMD database combination was used for mapping (Fig. 3A; Fig. 4). In addition, most of the dominant (relative abundance > 1%) and intermediate (relative abundance between 0.01% and 1%) fungal taxa were accurately (82.9%) assigned across the genus (Fig. 3A) and higher taxonomic levels (Fig. 4) if multiple HMDs combination was applied. Notably, with relative abundance greater than 0.01 %, the true positive rate (89.4%) was much higher than the false positive rate (10.6%) for fungal genera (Fig. 3A), as well as higher taxonomic levels (Fig. 4), indicating that MicroFisher can precisely predict dominant community composition. The absolute error identifies the percentage difference between the measured and actual abundance of each taxon was also calculated to evaluate the accuracy of the relative abundance prediction (Fig. 3B, Fig. S5). Overall, the relative abundance of fungal taxa estimated by MicroFisher using multiple HMDs has higher accuracy than single HMDs across genus (Fig. 3B) and other taxonomic levels (Fig. S5). At the genus level, the percentage of detected taxa with absolute abundance errors higher than 2.5% were lower for multiple HMDs dataset, ranging from 1.2% to 2.7%. On the other hand, they ranged between 5.4% and 6.9% for four different single HMDs. The results suggest this package can estimate the taxa abundance of the fungal community with a low absolute error rate across different taxonomic levels.

Similar patterns were found while evaluating the accuracy of predicted taxa abundance with r.m.s. errors. For example, the r.m.s. errors of 0.85, 0.82, and 0.68 were identified using ITS1+ITS2, LSU D1+D2, and ITS+LSU versus 1.42, 1.37, 1.27, and 1.22 were identified using ITS1, ITS2, LSU D1, and LSU D2, respectively (Fig. 3B; MiniHit length of 120 bp). Moreover, high correlations between actual and inferred relative abundance at genus (Pearson r = 0.87, p < 1e-100) and class (Pearson r = 0.99, p < 1e-100) levels were observed from the relative abundance table output while ITS1+ITS2 HMDs was applied for MicroFisher (Fig. 3C), as well as LSU D1+D2 and ITS+LSU (Fig. S6). Such high correlations between actual and inferred relative abundance were also observed across all taxonomic levels while multiple HMDs were applied (Fig. S7-S8). In addition, taxa with the real and inferred log-transformed abundance less than 5% showed high correlations (genus: r = 0.86, p < 1e-100; class: r = 0.91, p < 1e-50) (Fig. 3C, subfigure), suggesting that the MicroFisher can accurately predict the abundance of fungal taxa with relative abundances < 5%. Overall, the application of multiple HMDs combined with weight-based algorithms allows MicroFisher to sensitively and accurately classify the composition of dominant and intermediate fungal composition from a metagenome dataset.

### MicroFisher detects reliable fungal communities in real MG and MT data

We performed fungal community analysis for a set of soil MG and root eukaryotic MT data using MicroFisher in comparison with four other commonly used classification software (Eukdetect, Kaiju, Kraken2, and Metaphlan3). The default parameters of these tools were applied for the analysis. The combination of four HMDs (ITS1, ITS2, LSU D1, LSU D2) were used for MicroFisher workflow at MiniHit length of 80, 120, and 150 nt. The DNA amplicon sequencing data generated from the same soil was also included in the comparisons. In general, MicroFisher outperforms all other classification tools in fungal community profiling (Fig 5; Fig S9-S11).

**Fig. 5.**
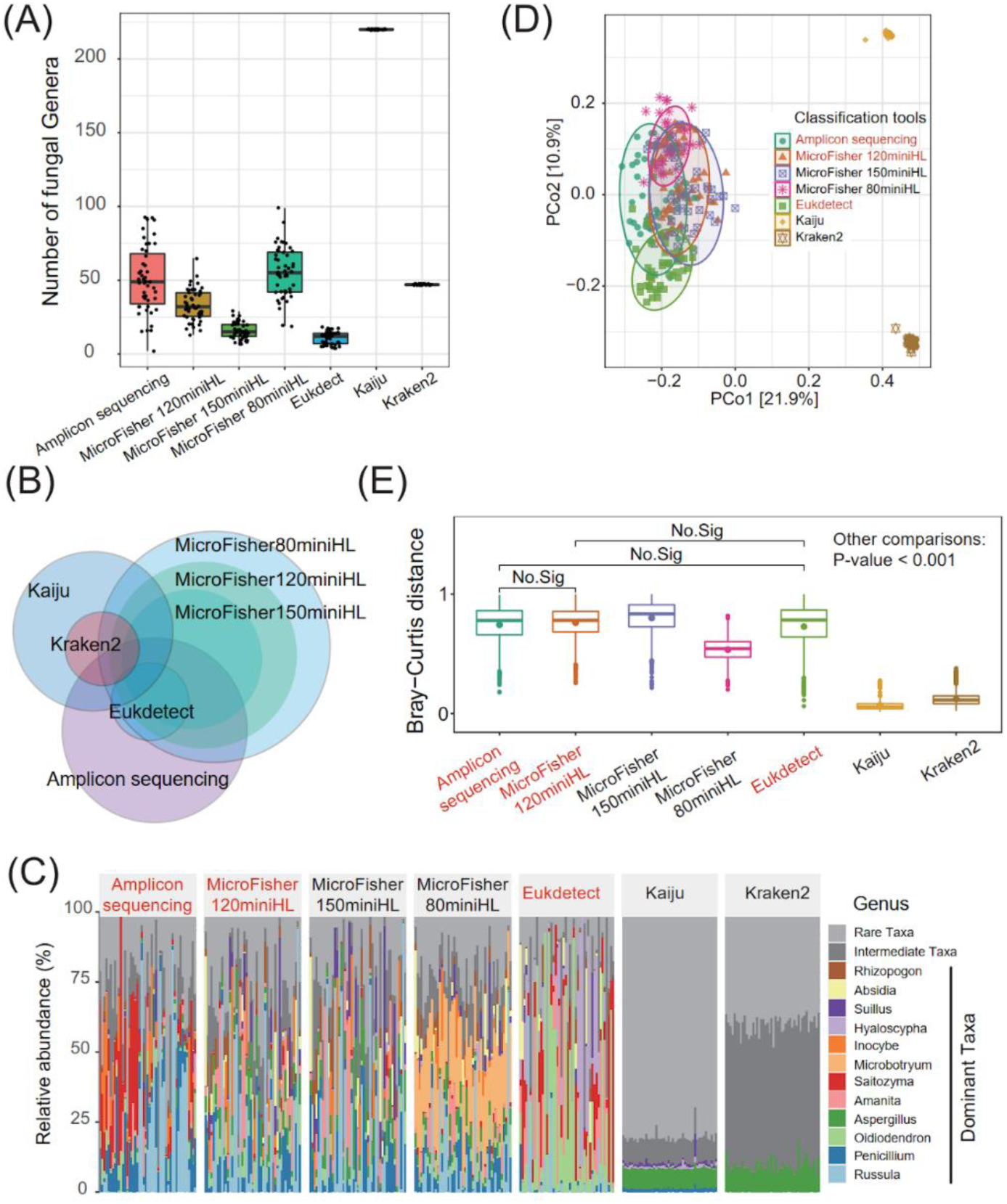
The analysis of soil metagenome data using MicroFisher versus other fungal classification tools. The DNA amplicon sequencing data generated from the same soil samples were also included in this comparison, indicating “amplicon sequencing”. Fungal profiling from amplicon sequencing was performed using QIIME2, and the metagenomic dataset was analyzed using MicroFisher with the minimum hit length of 80, 120, and 150 bp, Eukdetect, Kaiju, Kraken2, and Metaphlan3 (no fungal taxa detected, and is not shown) with default parameters. Only genus-level taxonomy was performed for this comparison. The analyses include: (A) Richness of soil fungal communities, and (B) Venn plot illustrates the overlap of fungal communities classified by different methods. (C) The dominant taxa (% abundance that > 1%) were obtained from different pipelines. (D) Principal Coordinates Analysis (PCoA) and (E) Bray-Curtis to identify the dissimilarity of soil fungal community composition resulting from different classification tools. MiniHL: minimum hit length; Dominant Taxa: the average relative abundance greater than 1%; Intermediate Taxa: the average relative abundance between 0.01% and 1%; Rare Taxa: the average relative abundance lower than 0.01%. MicroFisher classification and abundance evaluation were conducted using ITS+LSU HMDs with default parameters (MiniHit length of 120 bp, weighted and filter mode)

In the alpha-diversity estimation of MG data, on average, 50 genera were detected using amplicon sequencing across the soil samples, and 18, 31, and 56 genera were detected from soil MGs using MicroFisher at the MiniHit length of 150, 120, and 80 bp, respectively, while only 17 genera were recovered using Eukdetect (Fig. 5A). In addition, Metaphlan3 failed to detect fungal species from MGs and no significant difference (P > 0.05) between samples was found in the alpha diversity of the fungal community profiled by Kaiju and Kraken2. When comparing the shared genera detected from those MG data using selected classification tools to the amplicon sequencing result, Eukdetect and MicroFisher recovered more shared genera in comparison with Kraken2 and Kaiju (Fig. 5B). Also, MicroFisher recalled the highest number of shared genera (118 genera, MiniHit length of 120 bp) present in the amplicon sequencing result (Fig. 5B). Those results suggest that MicroFisher is more sensitive in fungal detection than the other examined classification tools. In addition, 177 and 185 genera were exclusively detected by amplicon sequencing and MicroFisher (MiniHit length of 120 bp), respectively. This might be due to the differences in sequencing technique, database application, and mapping algorithm utilized by these two approaches.

Similar patterns of relative abundance of dominant fungal taxa were observed using 120 bp and 150 bp MiniHit length compared to 80 bp MiniHit length in MicroFisher (Fig. 5C). This suggests that MiniHit length at 120 bp or longer provides more accurate outcomes in profiling fungal community abundance. Interestingly, the relative abundance of dominant fungal taxa was different among amplicon sequencing, Microfisher, and Eukdeteck. For example, relatively higher abundances of *Sailozyma*, *Russula*, and *Inocybe* were identified using DNA amplicon sequencing for many of the samples. While Microfisher detected *Russula*, *Suillus*, and *Amanita* with higher relative abundance. A relatively higher abundance of *Oidiodendron*, *Saitozyma*, and *Hyaloscypha* was detected in all of the samples using Eukdeteck. The abundance of overall dominant taxa was distributed more evenly across the samples using MicroFisher compared to Amplicon sequencing and Eukdetect. These examples indicate that compared to Eukdetect, the fungal community classified by amplicon sequencing and MicroFisher may be the more reliable tools in abundance estimation of the MG data. However, more studies will be needed to determine which of these two methods (amplicon sequencing or MicroFisher) can give more accurate results in the abundance estimation of fungi from soil DNA. In addition, the dominant taxa that were predicted by amplicon sequencing were not detected using Kaiju and Kraken2. Kaiju and Kraken2 failed to detect the difference in fungal community composition between soils collected from different field sites (Figs. 5C), suggesting the current protein/genome-based classification approaches may not perform well in the fungal classification of this MG dataset. This could be due to the lack of a robust genome database and the functional redundancy of protein databases used in these classification tools[7,8,47]. We also compared fungal beta-diversity classified using MicroFisher and other classification tools. Principal Coordinates Analysis (PCoA) and Bray-Curtis distance analysis indicated that the fungal communities classified by MicroFisher at MiniHit length of 120 nt and Eukdetect are more similar to the amplicon sequencing result (P-value > 0.05), reflecting MicroFisher may accurately predict the fungal composition and relative abundance at genus (Fig. 5D and 5E) and higher taxonomic level (Figs. S9D-E). This could also reflect that the marker gene prediction tool (Eukdetect) and ITS and LSU based prediction tools (Amplicon sequencing and MicroFisher) may give more similar outcomes in fungal community prediction compared with genome and protein prediction tools.

In the case of eukaryotic MT data generated from a root bioassay, 21, 11, and 5 genera on average were detected using MicroFisher at the MiniHit length of 80, 120, and 150 bp, respectively, while 7 genera on average were found by Eukdetect (Fig. S10A). Kaiju and Kraken2 predicted 221 and 48 genera, respectively, without significant differences between samples (P > 0.05). Although the number of fungal genera profiled by MicroFisher and Eukdetect are similar (Fig. S10A), the dominant genera identified using these two tools were mostly different (Fig. S10B). For example, using MicroFisher at MiniHit length of 120 bp, the dominant taxa identified include *Rhizopogon, Tomentella, Suillus,* and *Microbotryum*; while the fungal genera *Rhizopogon, Phialocephala, Rhizophagus, and Dactylonectria* were the predominant taxa detected by Eukdetect. Previous studies showed that the roots of pine seedlings grown in pine forest soil were typically dominated by a few specific ectomycorrhizal fungi, such as *Rhizopogon, Suillus, Peziza,* and *Tuber*[48–50], which is consistent with the classification results from MicroFisher at MiniHit length of 120 and 150 bp. However, Eukdetect and Kaiju failed to identify ectomycorrhizal fungi as the dominant taxa in around one-third of the root samples, and Kraken2 failed to identify the dominant ectomycorrhizal fungi across all the root samples (Fig. S10B), despite the fact that the root samples for MT analysis had a 68 ± 22 % colonization rate by ectomycorrhizal fungi. Beta-diversity comparison suggested that fungal communities classified by MicroFisher (MiniHit length of 120 and 150 bp) and Eukdetect have significant differences in composition compared with classification results from Kaiju and Kraken2 (P < 0.01) (Fig. S10 C and D). The class level comparison was also applied for the same MT dataset using these classification tools and received similar results as genus level comparison (Fig. S11), indicating the MicroFisher performs well in fungal classification across different taxonomic levels for MT data.

Overall, MicroFisher detected 306 (including 86 genera from 12 classes) and 142 clades (including 35 genera from 10 classes) with a relative abundance greater than 0.1 % in soil MGs and root MTs samples, respectively. MicroFisher also identified commonly found plant root symbiotic fungal taxa, including *Russula*, *Amanita*, *Inocybe*, *Suillus*, *Microbotryum*, *Wilcoxina*, *Sebacina*, *Trichoderma*, and *Tomentella*, in the pine forest system (Figs. 5F and S10B), which indicated that our approach was able to successfully recover the expected fungal taxa in certain small-scale studies. Thus, we highlight that MicroFisher outperforms other whole genome-based methods in fungal prediction, as the number of sequences and coverage of taxonomy in MicroFisher’s hypervariable marker databases are not limited by the size and by the targeted regions/genes of the database. Our results pave the way for the application of multiple hypervariable markers in profiling the composition and abundance of microbiomes from high-throughput sequencing.

## Discussion

With the development of high-throughput sequencing (such as MGs and MTs) that provides a better understanding of the relationships between population composition and ecological function, the impacts of fungal diversity and composition on ecosystem function and service have been well recognized. However, inferring microbial composition and abundance remains a significant challenge, as genome-reliant microbial classification often results in a high number of false-positive and missing predictions, especially when profiling fungal communities from short-reads.

To address this issue, we developed MicroFisher, which employs well-trained multiple hypervariable markers to classify fungal composition and estimate taxa abundance from metagenomic and metatranscriptomic sequencing data. Applying the well-trained multiple hypervariable marker databases is likely to have important implications for the accuracy of fungal classification and biological interpretation of ecological differences. The hypervariable marker databases comprise highly species-discriminative sequences to identify the fungal taxonomy from short-read (151 bp) sequencing. When conspired with the MicroFisher pipeline, which compiles the mapping-based classification and weight-based integration algorithms, multiple hypervariable markers show high sensitivity on both low– and high-complexity fungal community profiling. We also demonstrate that MicroFisher has substantially high accuracy and precision on fungal classification for fungal lineage at genus and higher taxonomic levels. To optimize the prediction performance of MicroFisher in fungal community profiling, the minimum hit length of 120 bp in fungal classification performs good in both sensitivity and accuracy.

Additionally, MicroFisher is substantially accurate in the relative abundance estimation of fungal communities. We highlight that MicroFisher outperforms alternative whole genome-based methods in fungal prediction, as the number of sequences and coverage of taxonomy in MicroFisher’s hypervariable marker databases are unrestricted by the limitations imposed by database size and the genome regions or genes. Our results pave the way for utilizing multiple hypervariable markers to characterize the composition and abundance of microbiomes from next generation sequencing.

In conclusion, we developed MicroFisher, an accurate and sensitive tool for fungal classification from metagenomic and metatranscriptomic datasets. By applying high-resolution hypervariable marker gene databases and weight-based integration algorithms, MicroFisher outperforms existing state-of-the-art tools. MicroFisher also enables the detection of rare taxa which is difficult to identify using other tools. This highlights the advancement of predicting fungal communities using multiple short hypervariable makers. Our results demonstrate the efficiency of MicroFisher for fungal classification and its potential for advancing the field of taxonomic profiling from metagenome and metatranscriptome.

## Data availability

All data, codes, and scripts used for this study are available online or upon request to the authors. The hypervariable marker databases generated in our study are available online at https://figshare.com/articles/dataset/MicroFisher_DBs/19679595; the MicroFisher is written in Python, and is available for download at https://github.com/NFREC-Liao-Lab/MicroFisher-Fungal-Profiling. Synthetic metagenomic sequencing datasets of the fungal ITS and rRNA genes used in this study are available at https://doi.org/10.6084/m9.figshare.20001542.v1. Metagenomic and metatranscriptomic sequencing data of the pine forest soils are available at the Joint Genome Institute (JGI) under Award DOI:10.46936/10.25585/60001406, the amplicon sequencing datasets of soil samples are available upon request to the authors. All codes, shell scripts, and R scripts are freely available at https://github.com/NFREC-Liao-Lab/Fisher-fungal-classification. The full tables of results for the fungal community classification and performance evaluation of simulated datasets using MicroFisher, and profiling results from real metagenomic and metatranscriptomic datasets using MicroFisher and other existing methods are available at Supplementary tables.

## Acknowledgment

This work was supported by several fundings to H-L L, including DE-SC0020403, DE-SC0012704, NSF IOS-PBI (2029168), USDA-NIFA (2019–67013-29107), USDA-NIFA Research Capacity Fund McIntire-Stennis accession no. 1026825 and Hatch accession no. 7001162. A portion of this research was performed on a project award JGI CSP (DOI: 10.46936/10.25585/60001406) under JGI’s community science program, and used resources at the DOE Joint Genome Institute (https://ror.org/04xm1d337), which are DOE Office of Science User Facility. The facility is sponsored by the Biological and Environmental Research program and operated under Contract Nos. DE-AC02-05CH11231 (JGI).

## Author contribution

H.L. Liao and R. Vilgalys coordinated the project; H. Wang and S. Wu developed the MicroFisher pipeline; H. Wang, K. Zhang and S. Wu performed the data analysis; K.H. Chen, R. Vilgalys, and H.L. Liao conducted the raw data generation for DNA amplicon, MG and MT data; H. Wang, K. Zhang and H.L. Liao wrote the manuscript; all authors contribute to manuscript edit.

## Figure legend

**Fig. S1** Representative sequence alignment example showing the sequence polymorphisms of LSU D1 and LSU D2 regions and the hypervariable regions. The multiple sequence alignment results with gaps were used to help identify the hypervariable regions, where the regions with low consensus were considered as the candidates of the “hypervariable region”, for example the “Hypervariable region 1” and “Hypervariable region 2”.

**Fig. S2** The sequence motifs illustrate the hypervariable regions in Internal transcribed spacer (ITS) and large subunit ribosomal RNA (LSU rRNA) genes. The hypervariable markers, including ITS1, ITS2, LSU D1, and LSU D2 were illustrated in sequence motifs (A and B). The Bits value represents the consensus of each base pair in the marker sequence. (C) Conceptual plots illustrate the structure and positions of hypervariable regions in ITS (above) and LSU rRNA D1D2 (below) sequences. The published primer sets (black arrows) were marked to show the approximate areas where the hypervariable regions were allocated.

**Fig. S3** Venn plot visualizes the overlap of fungal species in four generated hypervariable marker databases (HMDs).

**Fig. S4** The performance of MicroFisher’s fungal classification using single and multiple hypervariable marker databases (HMDs). The mock communities were simulated from randomly selected 50, 100, and 200 fungal species by InSilicoSeq, and the fungal classification and profiling were performed using MicroFisher (multiple HMDs applied) and Centrifuge (single HMD applied). The performance shows MicroFisher prediction at each taxonomy level by using a Minimum hit Length of 120 bp. The performance of accuracy, precision, and sensitivity were generated for each taxonomy level, the colors indicate the different number of species (50, 100, 200) used to generate the simulating datasets.

**Fig. S5** Absolute and root mean square (r.m.s) errors of predicted taxa abundance from 15 synthetic communities using single (ITS1, ITS2, LSU D1, and LSU D2; profiled by Centrifuge with minimum hit length of 120 bp) and multiple HMDs (ITS1+ITS2, LSU D1+D2, and ITS+LSU; profiled by MicroFisher with default setting). The taxa were ordered by value of absolute error and exhibited with the fraction of estimated taxa. Root mean square (r.m.s.) error was calculated for each mode, and the total number of truly predicted clades detected by different modes from 15 synthetic communities was shown as well. MicroFisher classification and abundance evaluation were conducted with default parameters (MiniHit length of 120 bp, weighted and filter mode).

**Fig. S6** Correlations of true and inferred taxa abundances for taxa estimated from 15 synthetic communities. Green and red dots denote taxa at the class and genus level respectively. To explore the correlation of taxa with abundance less than 5 %, we transformed them using log10 before performing the Pearson correlation. Pearson correlation coefficient and P-value for the taxa were calculated. MicroFisher classification and abundance evaluation were conducted using multiple HMDs: ITS+LSU (A) and LSU D1+D2 (B). The fungal classification was performed with default parameters (MiniHit length of 120 bp, weighted and filter mode).

**Fig. S7** Evaluation of Accuracy of relative abundance tables generated from 15 synthetic fungal community data using MicroFisher. Five taxonomy levels, including class (A), order (B), family (C), genus (D), and species (E), are reported here with the absolute errors in abundance estimation and rooted mean squared (r.m.s.) errors (left) of the predict abundance from hypervariable marker databases ITS1+ITS2 (red), LSU rRNA D1+D2 (blue), and ITS+LSU (green). Meanwhile, the correlations between true and predicted abundance values were shown (right). The abundance values were log10 transformed. Colors illustrate the predicted abundances from different marker databases: red, ITS1+ITS2; purple, LSU D1+D2; and green, ITS+LSU. MicroFisher classification and abundance evaluation were conducted with default parameters (MiniHit length of 120 bp, weighted and filter mode).

**Fig. S8** Evaluation of Accuracy of relative abundance tables generated from the synthetic community Simulating_200species_1 data using MicroFisher. Five taxonomy levels, including class (A), order (B), family (C), genus (D), and species (E), are reported here with the absolute errors in abundance estimation and rooted mean squared (r.m.s.) errors (left) of the predict abundance from hypervariable marker databases ITS1+ITS2 (red), LSU rRNA D1+D2 (blue), and ITS+LSU (blue). Meanwhile, the correlations between true and predicted abundance values were shown (right). The abundance values were log10 transformed. Colors illustrate the predicted abundances from different marker databases: red, ITS1+ITS2; purple, LSU D1+D2; and green, ITS+LSU. MicroFisher classification and abundance evaluation were conducted with default parameters (MiniHit length of 120 bp, weighted and filter mode).

**Fig. S9** The analysis of soil metagenome data using MicroFisher versus other fungal classification tools. The DNA amplicon sequencing data generated from the same soil samples were also included in this comparison, indicating “amplicon sequencing”. Fungal profiling from amplicon sequencing was performed using QIIME2, and the metagenomic dataset was analyzed using MicroFisher with the minimum hit length of 80, 120, and 150 bp, Eukdect, Kaiju, Kraken2, and Metaphlan3 (no fungal taxa detected, and is not shown) with default parameters. Class-level taxonomy was performed for this comparison. The analyses include: (A) Richness of soil fungal communities (B) Venn plot illustrates the overlap of fungal communities classified by different methods. (C) The dominant taxa (% abundance that > 1%) were obtained from different pipelines. (D) Principal Coordinates Analysis (PCoA) and (E) Bray-Curtis to identify the dissimilarity of soil fungal community composition resulting from different classification tools. MiniHL: minimum hit length; Dominant Taxa: the average relative abundance greater than 1%; Intermediate Taxa: the average relative abundance between 0.01% and 1%; Rare Taxa: the average relative abundance lower than 0.01%. MicroFisher classification and abundance evaluation were conducted using ITS+LSU HMDs with default parameters (MiniHit length of 120 bp, weighted and filter mode).

**Fig. S10** The analysis of root metatranscriptomic data using MicroFisher versus other fungal classification tools. Fungal profiling from the metatranscriptomic dataset was analyzed using MicroFisher with the minimum hit length of 80, 120, and 150 bp, Eukdect, Kaiju, Kraken2, and Metaphlan3 (no fungal taxa detected, and is not shown) with default parameters. Only genus-level taxonomy was performed for this comparison. The analyses include: (A) Richness of soil fungal communities (B) The dominant taxa (% abundance that > 1%) were obtained from different pipelines. (C) Principal Coordinates Analysis (PCoA) and (D) Bray-Curtis to identify the dissimilarity of soil fungal community composition resulting from different classification tools. MiniHL: minimum hit length; Dominant Taxa: the average relative abundance greater than 1%; Intermediate Taxa: the average relative abundance between 0.01% and 1%; Rare Taxa: the average relative abundance lower than 0.01%. MicroFisher classification and abundance evaluation were conducted using ITS+LSU HMDs with default parameters (MiniHit length of 120 bp, weighted and filter mode).

**Fig. S11** The analysis of root metatranscriptomic data using MicroFisher versus other fungal classification tools. Fungal profiling from the metatranscriptomic dataset was analyzed using MicroFisher with the minimum hit length of 80, 120, and 150 bp, Eukdect, Kaiju, Kraken2, and Metaphlan3 (no fungal taxa detected, and is not shown) with default parameters. Class-level taxonomy was performed for this comparison. The analyses include: (A) Shannon index of soil fungal communities (B) The dominant taxa (% abundance that > 1%) were obtained from different pipelines. (C) Principal Coordinates Analysis (PCoA) and (D) Bray-Curtis to identify the dissimilarity of soil fungal community composition resulting from different classification tools. MiniHL: minimum hit length; Dominant Taxa: the average relative abundance greater than 1%; Intermediate Taxa: the average relative abundance between 0.01% and 1%; Rare Taxa: the average relative abundance lower than 0.01%. MicroFisher classification and abundance evaluation were conducted using ITS+LSU HMDs with default parameters (MiniHit length of 120 bp, weighted and filter mode).

