## Supplemental figures and tables for "MicroFisher: Fungal taxonomic classification for metatranscriptomic and metagenomic data using multiple short hypervariable markers"

**Affiliations:**

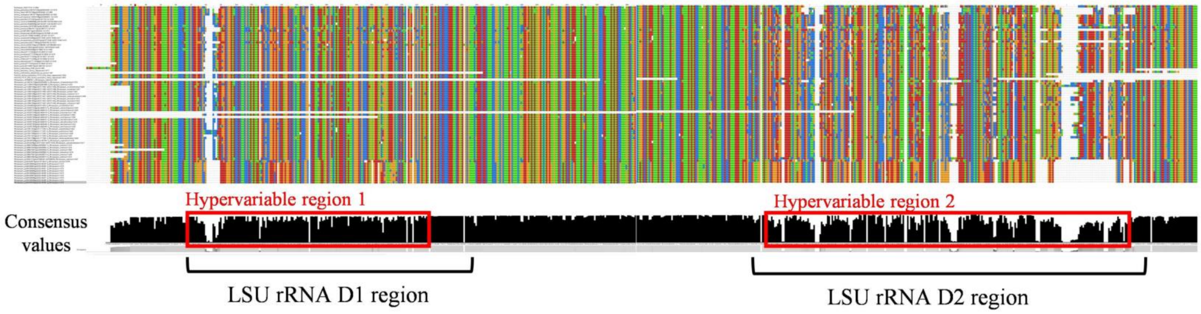


**Fig. S1** Representative sequence alignment example showing the sequence polymorphisms of LSU D1 and LSU D2 regions and the hypervariable regions. The multiple sequence alignment results with gaps were used to help identify the hypervariable regions, where the regions with low consensus were considered as the candidates of the "hypervariable region", for example the "Hypervariable region 1" and "Hypervariable region 2".

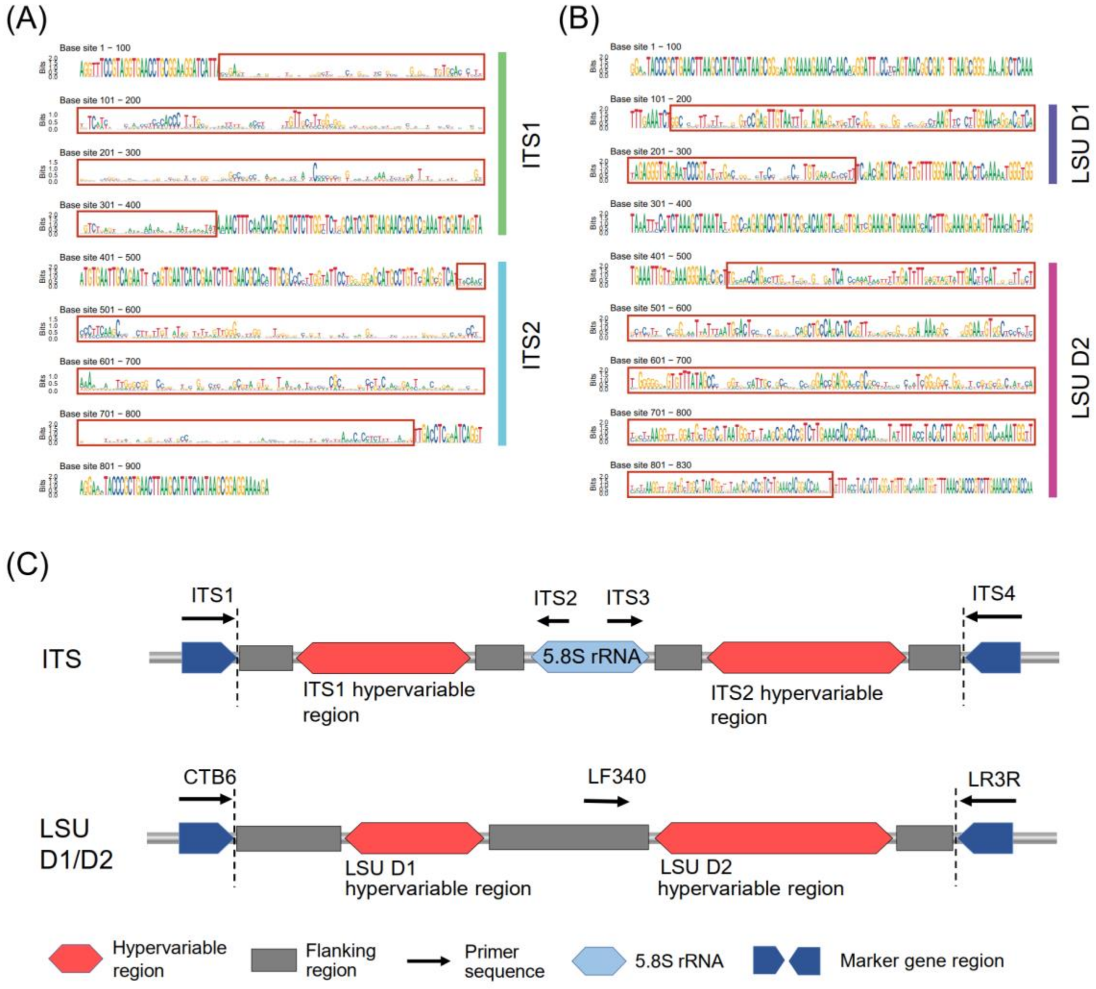


**Fig. S2** The sequence motifs illustrate the hypervariable regions in Internal transcribed spacer (ITS) and large subunit ribosomal RNA (LSU rRNA) genes. The hypervariable markers, including ITS1, ITS2, LSU D1, and LSU D2 were illustrated in sequence motifs (A and B). The Bits value represents the consensus of each base pair in the marker sequence. (C) Conceptual plots illustrate the structure and positions of hypervariable regions in ITS (above) and LSU rRNA D1D2 (below) sequences. The published primer sets (black arrows) were marked to show the approximate areas where the hypervariable regions were allocated.

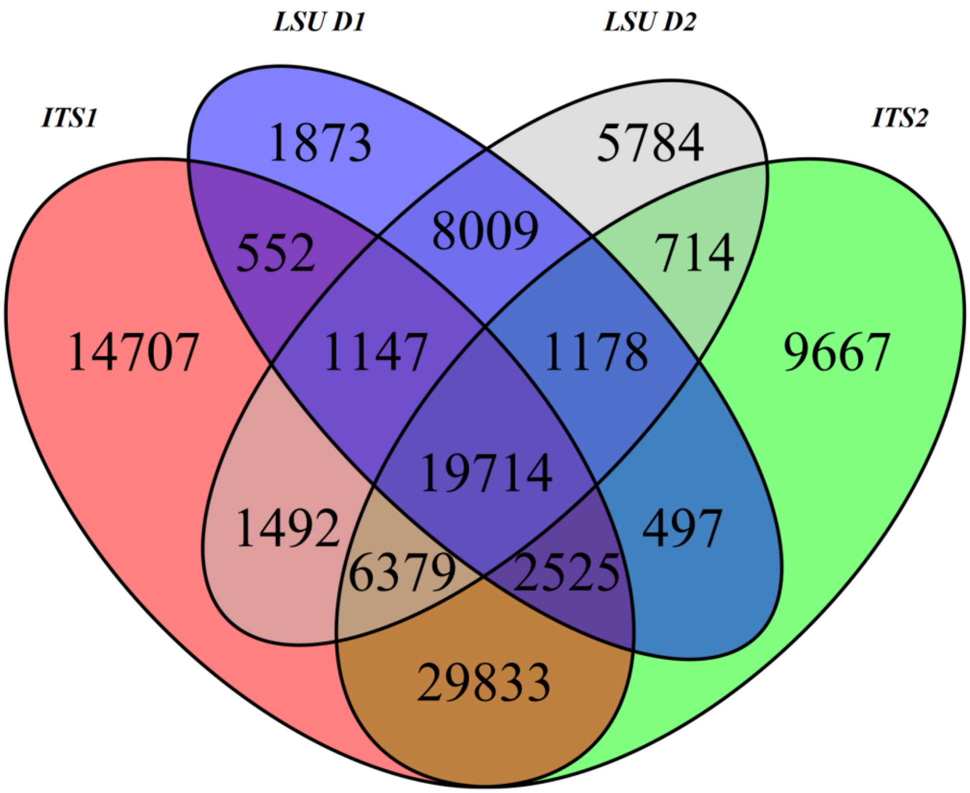


**Fig. S3** Venn plot visualizes the overlap of fungal species in four generated hypervariable marker databases (HMDs).

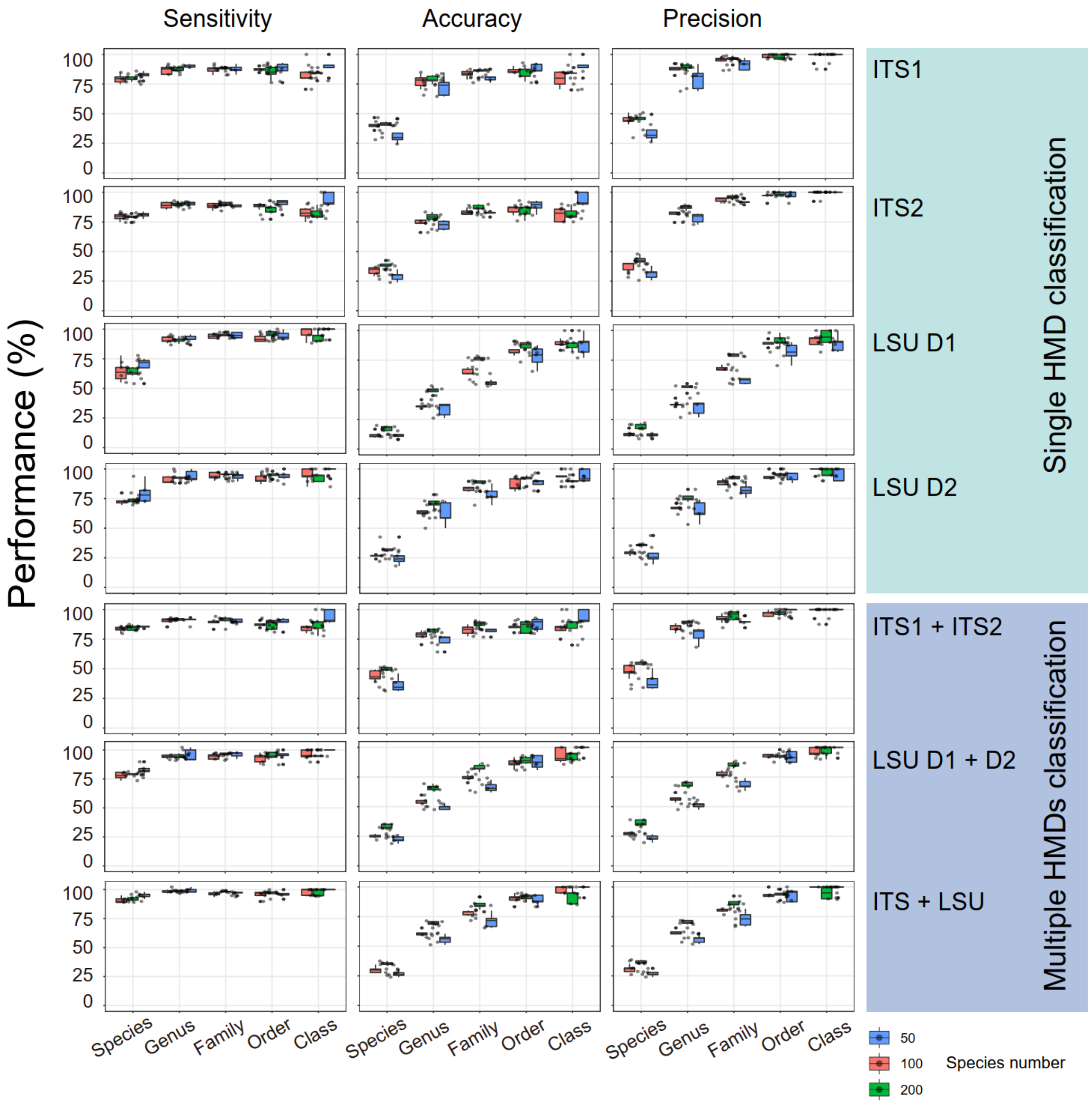


**Fig. S4** The performance of MicroFisher's fungal classification using single and multiple hypervariable marker databases (HMDs). The mock communities were simulated from randomly selected 50, 100, and 200 fungal species by InSilicoSeq, and the fungal classification and profiling were performed using MicroFisher (multiple HMDs applied) and Centrifuge (single HMD applied). The performance shows MicroFisher prediction at each taxonomy level by using a Minimum hit Length of 120 bp. The performance of accuracy, precision, and sensitivity were generated for each taxonomy level, the colors indicate the different number of species (50, 100, 200) used to generate the simulating datasets.

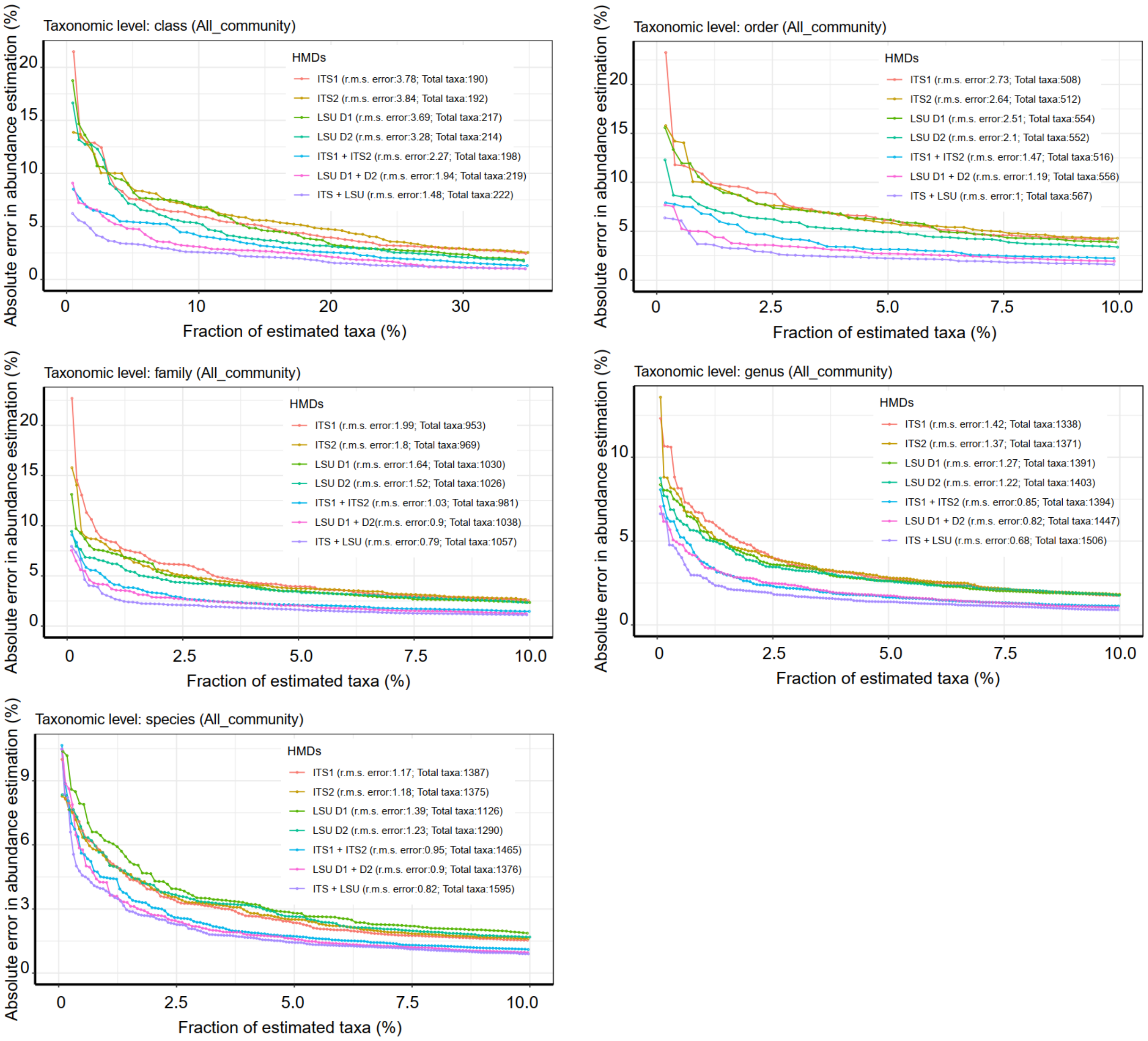


**Fig. S5** Absolute and root mean square (r.m.s) errors of predicted taxa abundance from 15 synthetic communities using single (ITS1, ITS2, LSU D1, and LSU D2; profiled by Centrifuge with minimum hit length of 120 bp) and multiple HMDs (ITS1+ITS2, LSU D1+D2, and ITS+LSU; profiled by MicroFisher with default setting). The taxa were ordered by value of absolute error and exhibited with the fraction of estimated taxa. Root mean square (r.m.s.) error was calculated for each mode, and the total number of truly predicted clades detected by different modes from 15 synthetic communities was shown as well. MicroFisher classification and abundance evaluation were conducted with default parameters (MiniHit length of 120 bp, weighted and filter mode).

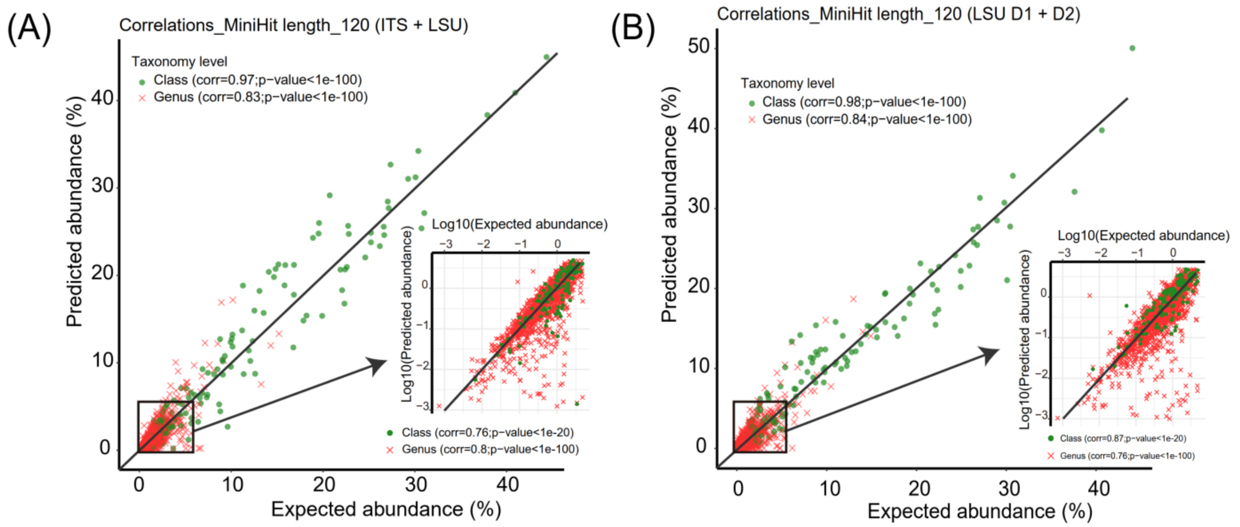


**Fig. S6** Correlations of true and inferred taxa abundances for taxa estimated from 15 synthetic communities. Green and red dots denote taxa at the class and genus level respectively. To explore the correlation of taxa with abundance less than 5 %, we transformed them using log10 before performing the Pearson correlation. Pearson correlation coefficient and P-value for the taxa were calculated. MicroFisher classification and abundance evaluation were conducted using multiple HMDs: ITS+LSU (A) and LSU D1+D2 (B). The fungal classification was performed with default parameters (MiniHit length of 120 bp, weighted and filter mode).

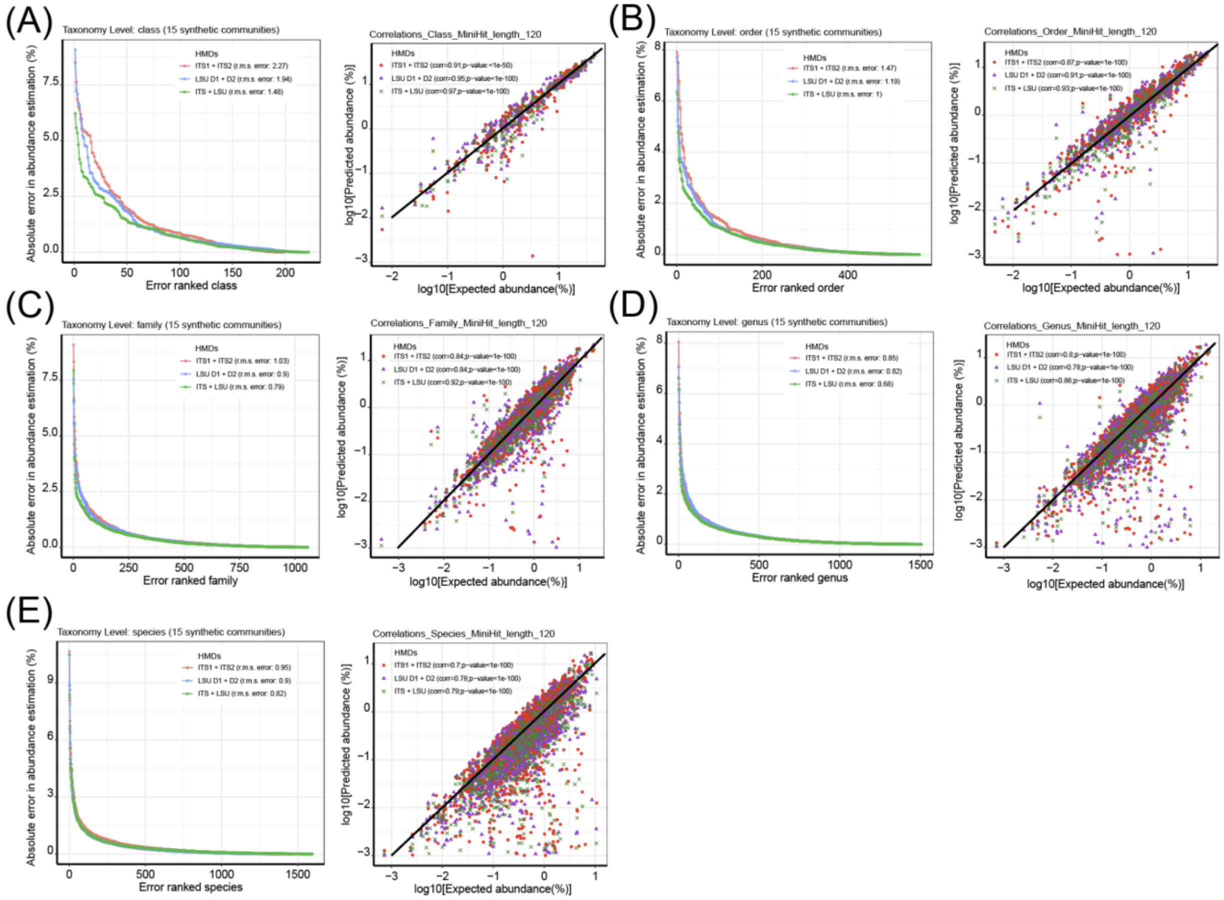


**Fig. S7** Evaluation of Accuracy of relative abundance tables generated from 15 synthetic fungal community data using MicroFisher. Five taxonomy levels, including class (A), order (B), family (C), genus (D), and species (E), are reported here with the absolute errors in abundance estimation and rooted mean squared (r.m.s.) errors (left) of the predict abundance from hypervariable marker databases ITS1+ITS2 (red), LSU rRNA D1+D2 (blue), and ITS+LSU (green). Meanwhile, the correlations between true and predicted abundance values were shown (right). The abundance values were log10 transformed. Colors illustrate the predicted abundances from different marker databases: red, ITS1+ITS2; purple, LSU D1+D2; and green, ITS+LSU. MicroFisher classification and abundance evaluation were conducted with default parameters (MiniHit length of 120 bp, weighted and filter mode).

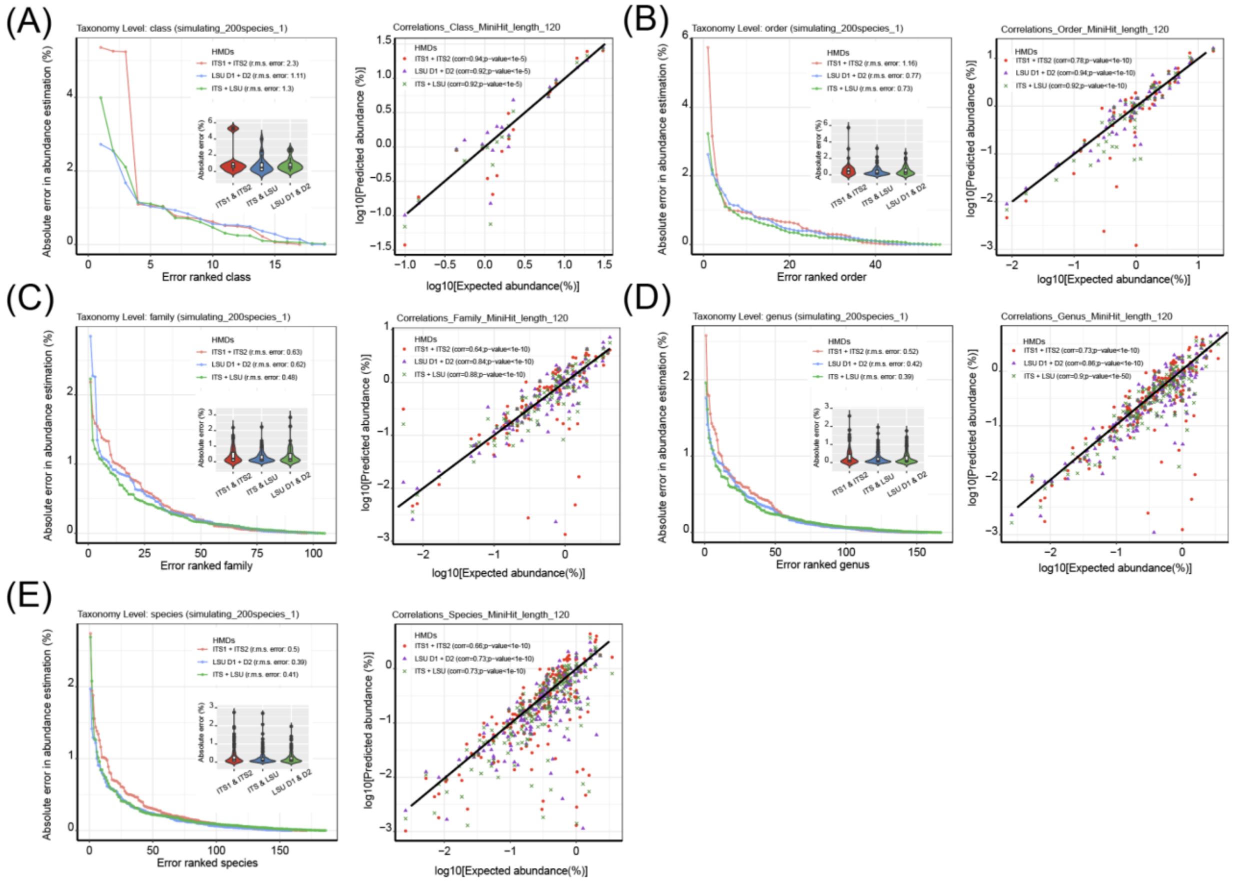


**Fig. S8** Evaluation of Accuracy of relative abundance tables generated from the synthetic community Simulating_200species_1 data using MicroFisher. Five taxonomy levels, including class (A), order (B), family (C), genus (D), and species (E), are reported here with the absolute errors in abundance estimation and rooted mean squared (r.m.s.) errors (left) of the predict abundance from hypervariable marker databases ITS1+ITS2 (red), LSU rRNA D1+D2 (blue), and ITS+LSU (blue). Meanwhile, the correlations between true and predicted abundance values were shown (right). The abundance values were log10 transformed. Colors illustrate the predicted abundances from different marker databases: red, ITS1+ITS2; purple, LSU D1+D2; and green, ITS+LSU. MicroFisher classification and abundance evaluation were conducted with default parameters (MiniHit length of 120 bp, weighted and filter mode).

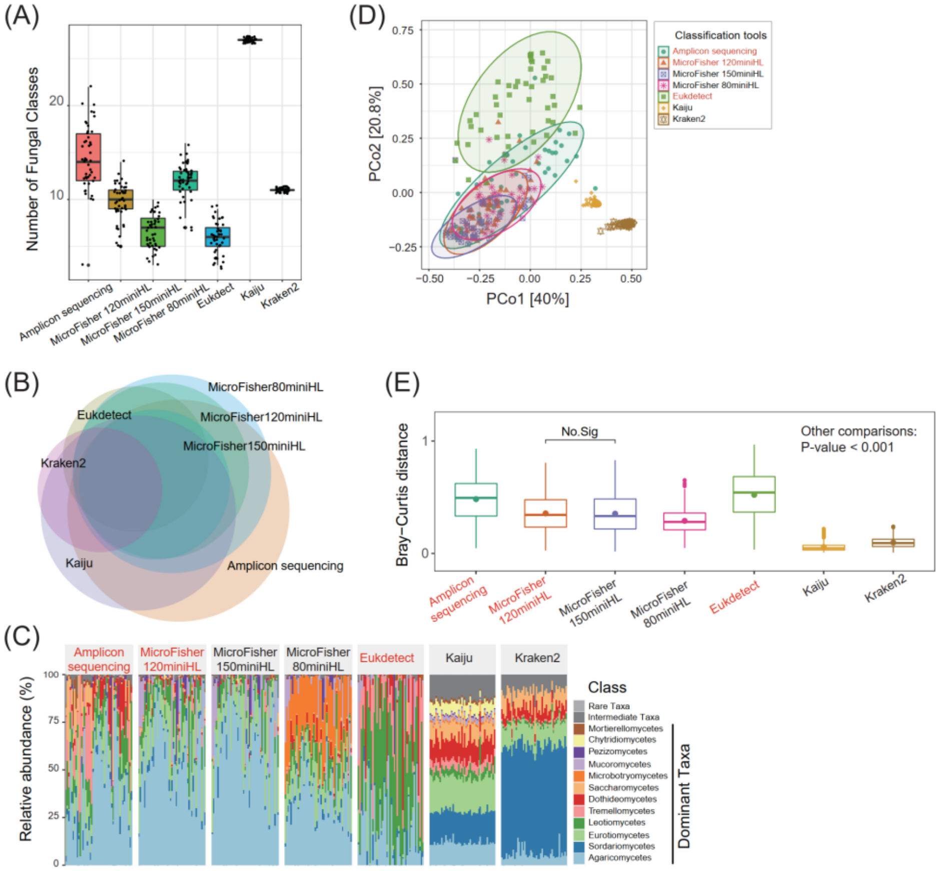


**Fig. S9** The analysis of soil metagenome data using MicroFisher versus other fungal classification tools. The DNA amplicon sequencing data generated from the same soil samples were also included in this comparison, indicating “amplicon sequencing”. Fungal profiling from amplicon sequencing was performed using QIIME2, and the metagenomic dataset was analyzed using MicroFisher with the minimum hit length of 80, 120, and 150 bp, Eukdect, Kaiju, Kraken2, and Metaphlan3 (no fungal taxa detected, and is not shown) with default parameters. Class-level taxonomy was performed for this comparison. The analyses include: (A) Richness of soil fungal communities (B) Venn plot illustrates the overlap of fungal communities classified by different methods. (C) The dominant taxa (% abundance that > 1%) were obtained from different pipelines. (D) Principal Coordinates Analysis (PCoA) and (E) Bray-Curtis to identify the dissimilarity of soil fungal community composition resulting from different classification tools. MiniHL: minimum hit length; Dominant Taxa: the average relative abundance greater than 1%; Intermediate Taxa: the average relative abundance between 0.01% and 1%; Rare Taxa: the average relative abundance lower than 0.01%. MicroFisher classification and abundance evaluation were conducted using ITS+LSU HMDs with default parameters (MiniHit length of 120 bp, weighted and filter mode).

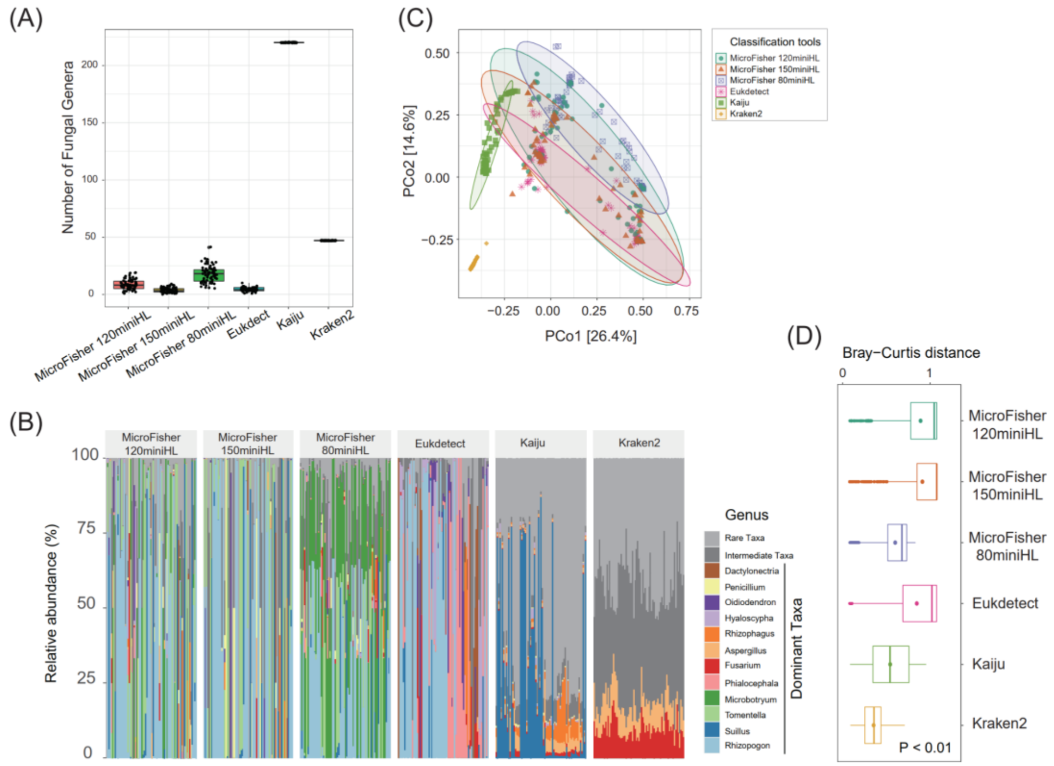


**Fig. S10** The analysis of root metatranscriptomic data using MicroFisher versus other fungal classification tools. Fungal profiling from the metatranscriptomic dataset was analyzed using MicroFisher with the minimum hit length of 80, 120, and 150 bp, Eukdect, Kaiju, Kraken2, and Metaphlan3 (no fungal taxa detected, and is not shown) with default parameters. Only genus-level taxonomy was performed for this comparison. The analyses include: (A) Richness of soil fungal communities (B) The dominant taxa (% abundance that > 1%) were obtained from different pipelines. (C) Principal Coordinates Analysis (PCoA) and (D) Bray-Curtis to identify the dissimilarity of soil fungal community composition resulting from different classification tools. MiniHL: minimum hit length; Dominant Taxa: the average relative abundance greater than 1%; Intermediate Taxa: the average relative abundance between 0.01% and 1%; Rare Taxa: the average relative abundance lower than 0.01%. MicroFisher classification and abundance evaluation were conducted using ITS+LSU HMDs with default parameters (MiniHit length of 120 bp, weighted and filter mode).

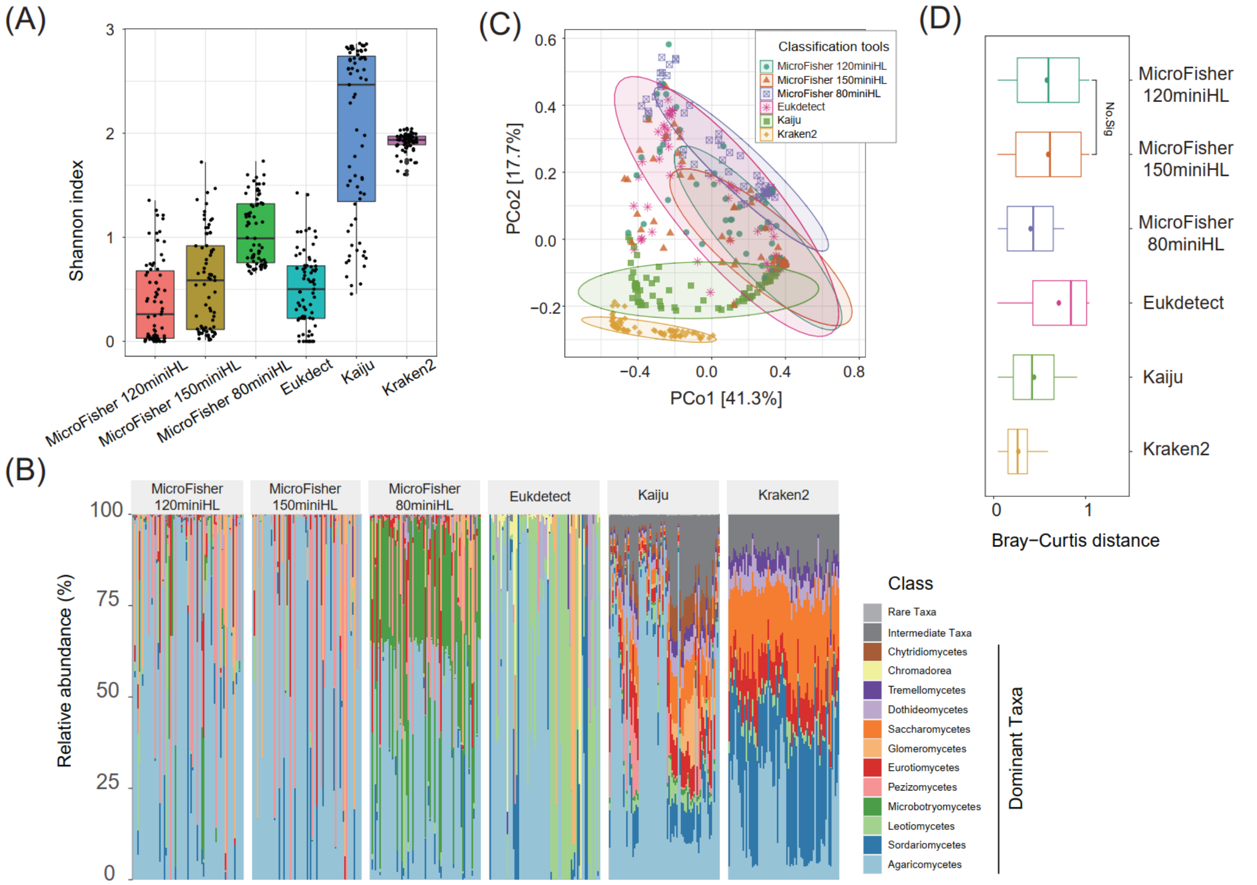


**Fig. S11** The analysis of root metatranscriptomic data using MicroFisher versus other fungal classification tools. Fungal profiling from the metatranscriptomic dataset was analyzed using MicroFisher with the minimum hit length of 80, 120, and 150 bp, Eukdect, Kaiju, Kraken2, and Metaphlan3 (no fungal taxa detected, and is not shown) with default parameters. Class-level taxonomy was performed for this comparison. The analyses include: (A) Shannon index of soil fungal communities (B) The dominant taxa (% abundance that > 1%) were obtained from different pipelines. (C) Principal Coordinates Analysis (PCoA) and (D) Bray-Curtis to identify the dissimilarity of soil fungal community composition resulting from different classification tools. MiniHL: minimum hit length; Dominant Taxa: the average relative abundance greater than 1%; Intermediate Taxa: the average relative abundance between 0.01% and 1%; Rare Taxa: the average relative abundance lower than 0.01%. MicroFisher classification and abundance evaluation were conducted using ITS+LSU HMDs with default parameters (MiniHit length of 120 bp, weighted and filter mode).

Table S1. Information of hypervariable marker databases provided by MicroFisher.

| Hypervariable Database | Version | No. of Sequences | Phyla | Classes | Orders | Family | Genera | Species | Average length (bp) | Sequences/ species |
|---|---|---|---|---|---|---|---|---|---|---|
| ITS1 | 1.0 | 405,672 | 9 | 58 | 219 | 752 | 5,416 | 76349 | 188.8 | 5.31 |
| ITS2 | 1.0 | 294,737 | 8 | 56 | 213 | 747 | 5428 | 70507 | 186.8 | 4.18 |
| LSU D1 | 1.0 | 81,704 | 8 | 59 | 225 | 799 | 5513 | 35495 | 172.8 | 2.30 |
| LSU D2 | 1.0 | 121,580 | 9 | 60 | 231 | 823 | 5927 | 44417 | 189.4 | 2.74 |
| Total | | 903,693 | 10 | 62 | 234 | 837 | 6545 | 104072 | | |

Table S2 The simulation datasets used for MicroFisher performance evaluation in this study.

| No. | Name | Organisms | Target genes | Sequences source | Number of species | Fungal richness |
|---|---|---|---|---|---|---|
| 1 | Simulating_200species_1 | Fungi | ITS, LSU | NCBI RefSeq database | 200 | High |
| 2 | Simulating_200species_2 | Fungi | ITS, LSU | NCBI RefSeq database | 200 | High |
| 3 | Simulating_200species_3 | Fungi | ITS, LSU | NCBI RefSeq database | 200 | High |
| 4 | Simulating_200species_4 | Fungi | ITS, LSU | NCBI RefSeq database | 200 | High |
| 5 | Simulating_200species_5 | Fungi | ITS, LSU | NCBI RefSeq database | 200 | High |
| 6 | Simulating_100species_1 | Fungi | ITS, LSU | NCBI RefSeq database | 100 | Middle |
| 7 | Simulating_100species_2 | Fungi | ITS, LSU | NCBI RefSeq database | 100 | Middle |
| 8 | Simulating_100species_3 | Fungi | ITS, LSU | NCBI RefSeq database | 100 | Middle |
| 9 | Simulating_100species_4 | Fungi | ITS, LSU | NCBI RefSeq database | 100 | Middle |
| 10 | Simulating_100species_5 | Fungi | ITS, LSU | NCBI RefSeq database | 100 | Middle |
| 11 | Simulating_50species_1 | Fungi | ITS, LSU | NCBI RefSeq database | 50 | Low |
| 12 | Simulating_50species_2 | Fungi | ITS, LSU | NCBI RefSeq database | 50 | Low |
| 13 | Simulating_50species_3 | Fungi | ITS, LSU | NCBI RefSeq database | 50 | Low |
| 14 | Simulating_50species_4 | Fungi | ITS, LSU | NCBI RefSeq database | 50 | Low |
| 15 | Simulating_50species_5 | Fungi | ITS, LSU | NCBI RefSeq database | 50 | Low |

Table S3 The real next-generation sequencing datasets used in this study.

| | Sequencing type | Sample source | Sample Type | Fungal richness | Sample number | Average reads number |
|---|---|---|---|---|---|---|
| 1 | Metagenome | Australia and the United States | Pine forest soil | High | 46 | ~500M |
| 2 | Metatranscriptome | Bioassay using soil from Australia and the United States | Bioassay pine seedling root | Middle | 71 | ~25M |
| 3 | Amplicon sequencing | Australia and the United States | Pine forest soil | High | 46 | ~100K |

Table S4. The glossary of terminology used in this paper

| Term | Abbreviation | Definition |
|---|---|---|
| Metagenomic | MG | Metagenomics is the study using genetic materials (genome sequences) recovered directly from environmental or clinical samples by a method called sequencing |
| Metatranscriptomic | MT | Metatranscriptomics is the analysis of the collective transcriptomes (sum total of all the messenger RNA molecules) of a given habitat. |
| Synthetic metagenomic | - | The synthetic metagenomic data that was simulated by InSilicoSeq. |
| Ribosomal RNA | rRNA | Ribosomal ribonucleic acid (rRNA) is a type of non-coding RNA that is the primary component of ribosomes, essential to all cells. |
| Small Subunit Ribosomal RNA | SSU | The small subunit ribosomal RNA (SSU rRNA) gene is the widely used molecular taxonomic marker for microorganisms |
| Large Subunit Ribosomal RNA | LSU | The large subunit ribosomal RNA (LSU rRNA) gene is the widely used molecular taxonomic marker for microorganisms |
| Internal Transcribed Spacer | ITS | The ITS regions span between the 18S rDNA and 5.8S rDNA (referred to as ITS1) and between the 5.8S rDNA and 28S rDNA (referred to as ITS2) in the eukaryotic genome and are highly variable. Hence, they provide sufficient information for the classification of eukaryotic microbes (fungi and yeasts) up to the species level. |

| | | |
|---|---|---|
| Inverse weighting | - | Inverse weighting is a statistical technique for calculating statistics standardized to a pseudo-population different from that in which the data was collected. |
| True positive | TP | A true positive is an outcome where the model correctly predicts the positive class. |
| False positive | FP | A false positive is an outcome where the model incorrectly predicts the positive class. |
| False negative | FN | A false negative is an outcome where the model incorrectly predicts the negative class. |
| Sensitivity (recall) | - | Recall is the ratio of the correctly labeled by our program to all who are expected.<br>Recall = TP/(TP+FN) |
| Accuracy | - | It's the ratio of the correctly labeled subjects to the whole pool of subjects.<br>Accuracy = (TP+TN)/(TP+FP+FN) |
| Precision | - | Precision is the ratio of the correctly labeled by our program to all labeled.<br>Precision = TP/(TP+FP) |
| Absolute error | - | Absolute error is the difference between measured or inferred value and the actual value of a quantity.<br>For example, if the real value of relative abundance is 35.67% while the predicted relative abundance is 35%, the absolute error (%) is 0.67 (%) |
| Root mean square error | r.m.s error/RMSE | Root Mean Square Error (r.m.s error, RMSE) is a standard way to measure the error of a model in predicting quantitative data. |

| | | |
|---|---|---|
| | | $$RMSE = \sqrt{\frac{\sum_{i=1}^{N} \lVert y(i) - \hat{y}(i) \rVert^2}{N}},$$ where N is the number of data points, y(i) is the i-th measurement, and $\hat{y}(i)$ is its corresponding prediction. |
